# Targeted over-expression of PPARγ in pig skeletal muscle improves oxidative fiber formation and intramuscular fat deposition

**DOI:** 10.1101/2020.06.24.168641

**Authors:** Hao Gu, Ying Zhou, Jinzeng Yang, Jianan Li, Yaxin Peng, Xia Zhang, Yiliang Miao, Wei Jiang, Guowei Bu, Liming Hou, Ting Li, Lin Zhang, Xiaoliang Xia, Zhiyuan Ma, Yuanzhu Xiong, Bo Zuo

## Abstract

PPARγ is a master regulator of adipogenesis and lipogenesis. To understand its roles in fat deposition and fiber formation in skeletal muscle, we successfully generated muscle-specific overexpression of PPARγ by pig cloning procedures. The content of intramuscular fat was significantly increased in PPARγ transgenic pigs while the muscle mass as lean percentage of body weight was not changed. The proteomics analysis demonstrated oxidative metabolism of fatty acids and respiratory chain were increased in PPARγ pigs. Furthermore, expressions of oxidative muscle fiber-related genes such as MyHC1 and TNNT1 were enhanced. CAMK2, MEF2 and PGC1α were also significantly increased in skeletal muscle of PPARγ pigs, indicating that Ca^2+^-sensitive phosphatases and kinases may play an important role in the switch of muscle fiber types when PPARγ activity is elevated in skeletal muscle. The results support skeletal muscle-specific overexpression of PPARγ can promote oxidative fiber formation and intramuscular fat deposition in pigs.

## Introduction

Peroxisome proliferator-activated receptor gamma (PPARγ) is a ligand-activated transcription factor from the nuclear receptor family of peroxisome proliferator-activated receptors (PPARs), it serves as the crucial regulator of adipogenesis, lipogenesis and lipid storage, and whole-body insulin sensitivity through coordinated activities in liver, muscle and adipose tissue (Evans et al., 2004). PPARγ is necessary for the formation and effectivity of fat cells (Lehrke et al., 2005; Mota et al., 2017). In mammals, PPARγ exists as two isoforms, PPARγ1 and PPARγ2. The expression of PPARγ protein in muscle is two-thirds of that present in adipose tissue, but PPARγ1 seems to be the predominant isoform expressed in skeletal muscle (Loviscach et al., 2000). PPARγ2 contains additional 30 amino acids at the N-terminal of PPARγ1, expressed mainly in adipose tissue (Mueller et al., 2002; Tontonoz et al., 1994; Zhu et al., 1995). During adipocyte differentiation, PPARγ is induced and forced expression in non-adipogenic cells which can effectively convert them into mature adipocytes (Tontonoz et al., 1994), while PPARγ knockout mice failed to develop adipose tissue (Barak et al., 1999; Kubota et al., 1999; Rosen et al., 1999). Studies have shown that when ectopic expression PPARγ and C/EBP alpha in myoblast cell culture, it induced myoblast cells into mature fat cells (Hu et al., 1995).

Muscle cells differ in rate of contraction and metabolic activities. Based on energy metabolism, muscle fibers can be divided into two types: slow/type I/oxidative fiber and fast/type II /glycolytic fiber. Type I muscle contains relatively high levels of mitochondria, supply the energy for the oxidation breathing (Nemeth et al., 1984), and have relatively high fat contents. Type I muscle has a slow ATPase rate and are adapted for long, slow contraction. Fast-twitch muscles (type II) express muscle myosin that has a fast ATPase rate and the cells tend to be larger. We previously demonstrated the proportion of Type I muscle was increased in skeletal muscle by over-expression PGC1 alpha in transgenic mice and pigs (Zhang et al., 2017). Increasing the number of type I muscle fibers will lead to more red color and flavor of meat. A high ratio of Type I muscle fibers is positively correlated with intramuscular fat content (Brocks et al., 1998; Renand et al., 2001).

Intramuscular fat (IMF) is located between and within muscle fibers, which is another important economic trait (i.e. marbling score) in pork production since phospholipids in IMF is the precursor of meat flavor (Wood et al., 2008). IMF content varies between pig breeds and between muscle types in the same breed. Variability in IMF content is mainly linked to the number and size of intramuscular adipocytes. Higher levels of IMF or marbling in pork can positively influence the juiciness, tenderness, and flavor of pork (Wood et al., 2004). It has been reported that pig breeds which have higher PPARγ2 expression level, have more IMF deposition (Grindflek et al., 1998). In our previous work, we found ectopic overexpression of swine PPARγ2 in skeletal muscle of mice can increase multiple gene expressions for adipocyte formation and elevated the triacylglycerol content in skeletal muscle (Huang et al., 2012).

In consideration of PPARγ’s important role in adipocyte cells and possible influence on the intramuscular fat deposition, we generated skeletal muscle-specific PPARγ overexpression transgenic pigs using CRISPR/Cas9-mediated site-specific integration and random integration methods. Both transgenic PPARγ pigs showed similar results that overexpress PPARγ in pig muscle can stimulate the muscle fiber conversion from fast to slow twitch and promote the deposition of intramuscular fat through neuromuscular Ca^2+^ signal regulation.

## Results

### 1. PPARγ transgenic pigs generated by random integration

In this work, somatic cell nuclear transfer (SCNT) and random integration technologies were used to generate transgenic pigs with muscle specific PPARγ overexpression. As shown in Fig. S1A, pN1-MCK-PPARγ vector containing 7kb full-length MCK promoter followed by PPARγ2’s cDNA sequence was constructed, and transfected into pig embryonic fibroblast cells from a 30-day-old Large White pigs. Then we selected a cell clone after G418 selection for SCNT (Fig. 1A). Embryos were transferred into 12 surrogate pigs. Eight pregnancies were established. And 19 piglets were born by natural delivery, of which 13 piglets are transgenic (TG) pigs according to PCR verification (Fig. 1B). After crossing with American wild-type (WT) Large White sows, we got 8 F1 TG pigs from one litter (Fig. S1B) and 9 F2 TG pigs from two litters (Fig. S1D). Southern blotting results showed the genetic stability of foreign PPARγ gene in the genomes of F0 and F1 transgenic pigs (Fig. 1C and S1C). Real-time PCR results in 11 different tissues of F1 pigs showed PPARγ expression was significantly increased in longissimus dorsi muscle, biceps femoris muscle, and gastrocnemius muscle of TG pigs (Fig. 1D). Western blotting results were also confirmed the higher PPARγ expression levels in longissimus dorsi muscle and gastrocnemius muscle of TG pigs (Fig. 1E). The meat trait measurements of F1 and F2 pigs showed that there was no significant difference for birth weight and lean meat percentage between TG and WT pigs, while IMF was significantly increased 0.8% and 0.7% in PPARγ F1 and F2 TG pigs separately (2.3±0.2% for F1 TG; 1.5±0.2% for F1 WT; P=0.016; 2.4±0.3% for F2 TG; 1.7±0.1% for F2 WT; P=0.008) (Fig. 1G, Table S1 and S2). H&E staining results further illustrated that there were more adipose cells and connective tissues in the longissimus dorsi muscle of TG pigs (Fig. 1F). These results showed the IMF content was significantly increased in PPARγ TG pigs without changing the lean meat percentage.

**Fig. 1.**
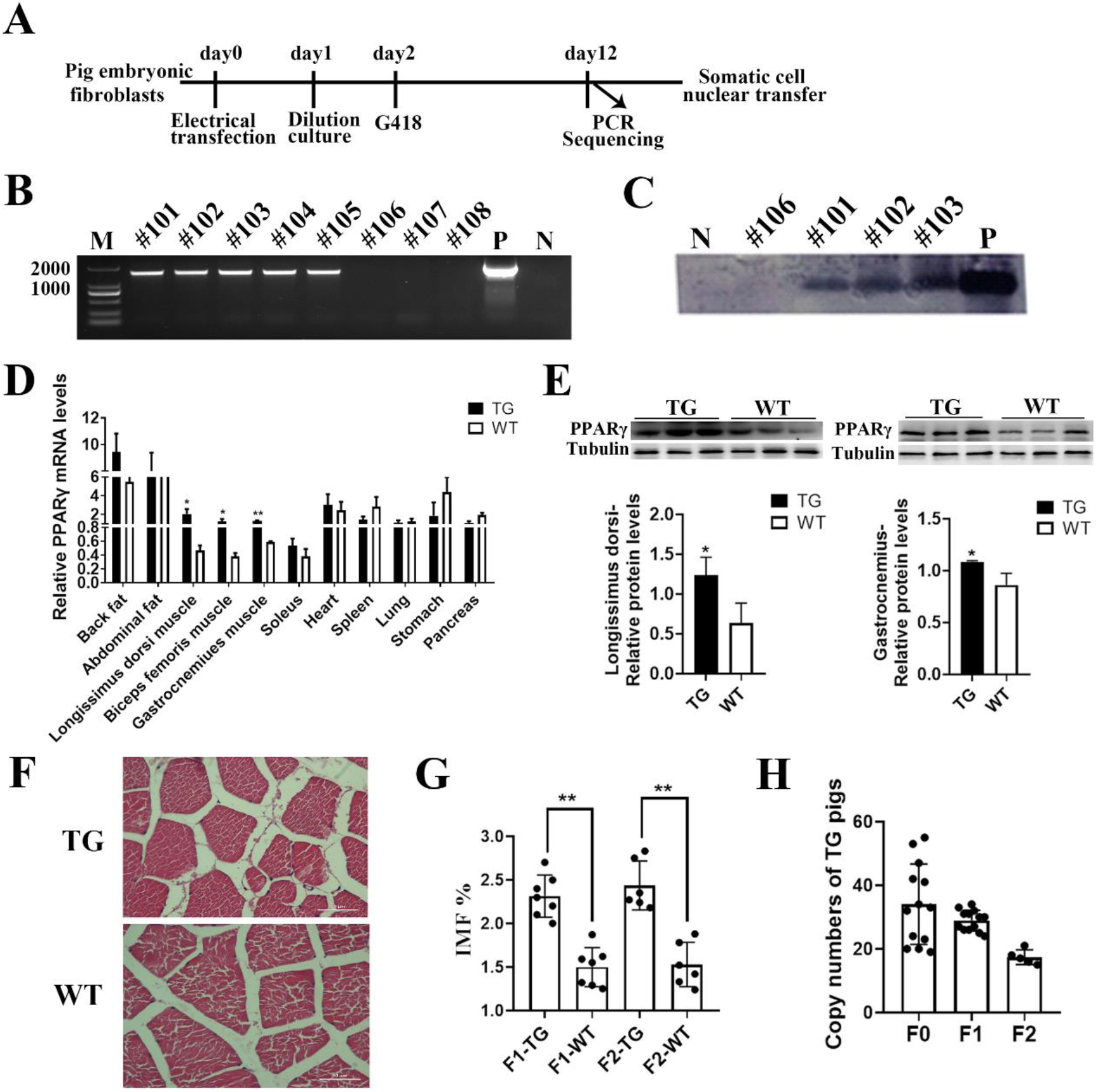
Muscle specific overexpression of PPARγ improves the IMF in PPARγ TG pigs. (A) Schematic illustration of the screening process to obtain TG monoclonal cell lines. (B, C) PCR identification (B) and Southern blotting (C) of F0 TG pigs. Lane N represents negative control from WT pigs; lane P represents positive control from vector. (D) The relative *PPARγ* mRNA expression levels in different tissues of F1 TG pigs compared with WT pigs (n=3). (E) Western blot analysis of PPARγ in muscles of TG pigs (n=3). (F) H&E staining on longissimus dorsi of TG and WT pigs. Scale bar: 50 μm. (G) Statistical results of slaughter indexes of F1 and F2 generation pigs’ intramuscular fat (IMF). (H) Statistical results of TG pigs’ copy numbers. The relative mRNA and protein levels were normalized to Tubulin. The data are presented as mean ± SD of independent experiments; * *p* < 0.05, ** *p* < 0.01. Dot plot reflects data points from independent experiment.

### 2. The generation of PPARγ KI pigs using CRISPR/Cas9-mediated site-specific integration in Rosa26 locus

When we test the copy number of PPARγ TG pigs, PPARγ copies were decreased in the F2 generation (Fig. 1H), indicating that the copy number of TG pigs by random integration was unstable. Therefore, we generated PPARγ knock-in (KI) pigs using SCNT with CRISPR/Cas9 system. The pig *Rosa26* locus which has been validated as the safe harbor (Li et al., 2014), was selected as the knock-in site. Approximately 1.0 kb upstream and 1.5 kb downstream of the cleavage site were used as the left homology arms (HA-L) and right homology arms (HA-R) for homologous recombination (HR), respectively. In the donor plasmid, the insertion sequence between HA-L and HA-R, contained an expression cassette consisting of porcine PPARγ2 cDNA driven by a shorter MCK promoter and a loxP-flanked positive selection cassette consisting of a neomycin resistance gene (neoR) expressed by the phosphoglycerate kinase promoter (PGKp) (Fig. 2B). Besides, the donor plasmid also carries a negative selection cassette in the non-insertion sequence, which contained a *HSV-TK* gene and a PGK promoter to inhibit random insertion.

**Fig. 2.**
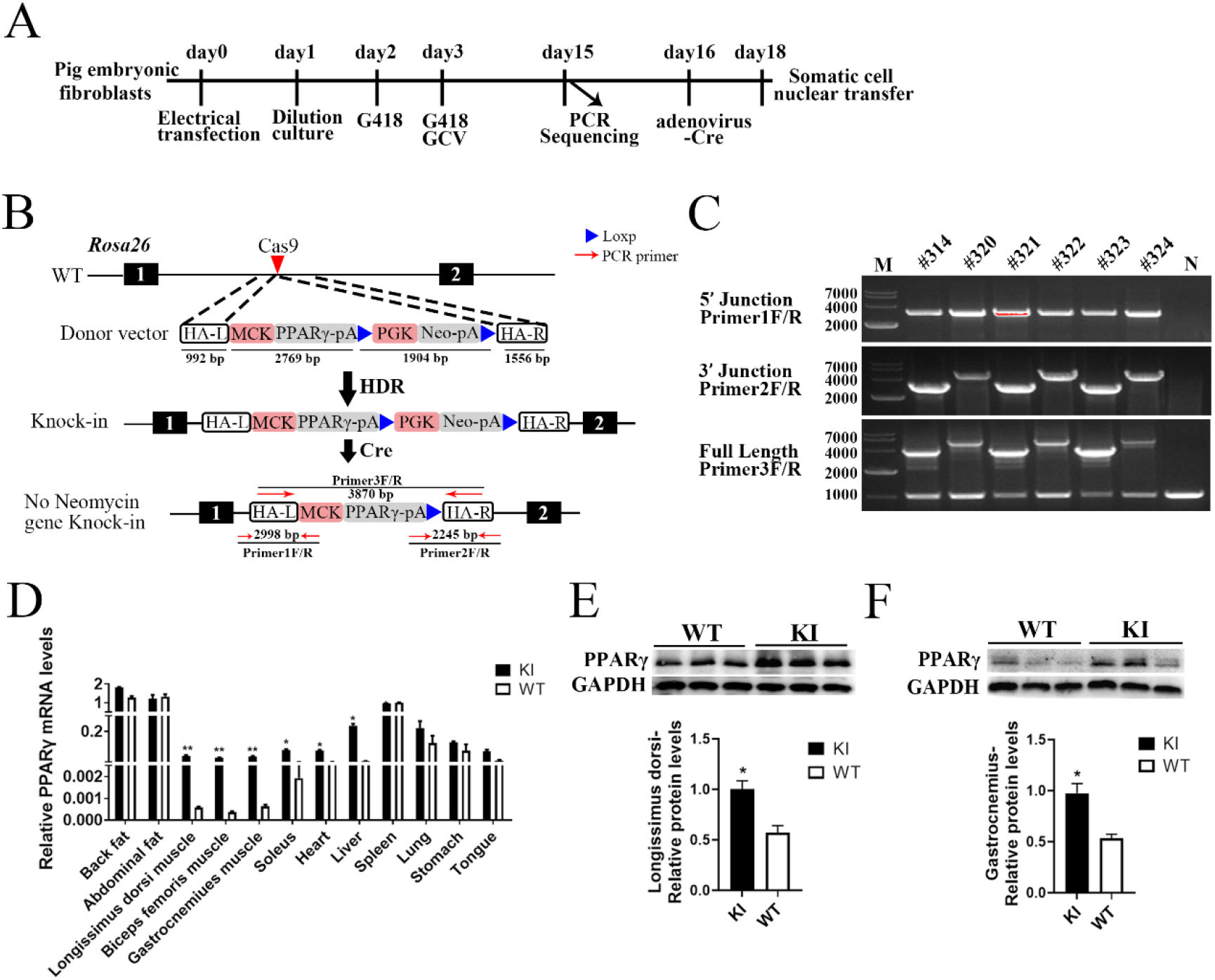
Generation of PPARγ KI pigs using CRISPR/Cas9. (A) Schematic illustration of the screening process to obtain monoclonal cell lines. (B) Schematic representation of the strategy to insert MCK-PPARγ cassette into the Rosa26 locus. Exons of Rosa26 are shown as black boxes, and the magenta arrowhead in the first intron represents the sgRNA targeting site. The donor vector which is flanked by the left and right homologous arms (HA-L and HA-R), contains the porcine PPARγ cDNA driven by Muscle Creatine Kinase (MCK) promoter and a loxP-flanked (blue arrowhead) neomycin expression cassette driven by the phosphoglycerate kinase promoter (PGK). The opposite red arrows in the bottom diagram indicate three pairs of primers designed for genotyping. Primer 1F/R for 5’junction PCR, primer 2F/R for 3’junction PCR and primer 3F/R for the full-length PCR. (C) The PCR verification of the site-specific targeting in the KI pigs. Lanes #314-#324 represent 6 founder (F0) male KI pigs, and lane N represent negative control WT pigs. (D) The relative PPARγ mRNA expression levels in different tissues of F1 KI pigs compared with WT pigs (n=3). (E, F) Relative PPARγ protein levels by Western blotting in the longissimus dorsi muscle (E) and gastrocnemius muscle (F) of KI pigs compared with WT pigs (n=3). The relative mRNA and protein levels were normalized to GAPDH. The data are presented as mean ± SD of independent experiments; * *p* < 0.05, ** *p* < 0.01.

The Cas9-sgRNA and donor plasmids were co-transfected into pig embryonic fibroblasts (PEFs) by electroporation, and then the cells were diluted into 96-well plates. G418 was added 2 days later, G418 and GCV were added 3 days later. After 12 days of continuous culture under positive (G418) and negative (GCV) selection, each single colony was detected by PCR and sequencing to select the KI cell lines. The transgenic cell lines were further cultured for 2 days in the medium supplemented with adenovirus-Cre to remove the neoR gene. The schematic for the production of KI pigs was shown in Fig. 2A. We selected the KI cell lines for SCNT and obtained 2075 embryos which were transferred into 10 surrogate pigs. Five pregnancies were established, and 13 piglets were born by natural delivery from four litters of which 6 were survived finally (Table S3). Genotyping by PCR analysis of 5’ junction and 3’ junction showed that all 6 male piglets have a correct site-specific insertion. Three of them (#314, #321, #323) were free of marker gene which means has no neomycin sequence at 3’ junction region. We found all the KI pigs were heterozygous when we proceeded the full-length PCR analysis, showing that the donor vectors were only inserted into one chromosome (Fig. 2C and Fig. S2 A). The full-length PCR
sequencing results of #314 KI pig were shown in Date S1.

The male founder (F0) pig (#314) was used to mate with two Danish WT Large White sows, 32 F1 pigs were generated. PCR results showed that there were 16 KI pigs and 16 WT pigs (Fig. S2 B), showing Mendelian segregation of transgenes among the F1 progeny (Table S4). We analyzed the copy number of pigs from each KI generation using real-time PCR, the results showed that the copy numbers were consistent among generations, that they all have one copy of insertion (Fig. S2 C). According to the location and sequence similarities of sgRNA, we used Cas-OFFinder online software to predict the potential off-target sites (Fig. S3 A). The PCR detection results revealed no off-targets in KI pigs (Fig. S3 B).

To verify muscle specific expression of the transgene, we tested PPARγ relative expression levels by real-time PCR in 12 different tissues from 5-month-old KI pigs and their WT siblings. As an endogenous gene, PPARγ has levels of expression especially in fat tissue (Fig. 1D). But with the MCK promoted overexpression of PPARγ, the PPARγ’s expressions were significantly increased in muscles at mRNA levels (Fig. 1D and S4 A) and also protein levels (Fig. 2E and 2F), which consistent with our random insertion results. The expression was also increased in the heart and liver by real-time PCR test, but no significant difference was observed on protein levels (Fig. S4 B). PPARγ’s expression pattern didn’t change in other tissues including backfat, abdominal fat, spleen, lung, stomach or tongue (Fig. 1D and S4 A). The morphology observations by H&E staining and weighting of heart, liver, spleen, lung, kidney and backfat also showed there were no significant difference between KI and WT pigs for these tissues (Fig. S5 A and S5 B).

### 3. PPARγ KI Pigs had higher IMF and enhanced adipogenesis related genes expressions

We recorded the pigs weight monthly, the growth curve showed there was no difference for birth weight and the growth rate between KI pigs and WT pigs (Fig. S6 A). Then we measured the carcass and meat quality traits of these pigs weighted about 100kg. The results showed that the IMF was increased by 0.47% in KI pigs (2.82±0.28% for KI; 2.35±0.16% for WT; P=0.0036), the meat marbling score was higher in KI pigs (2.86±0.35 for KI; 2.42±0.19 for WT; P=0.0136), while there was no significant difference in lean meat percentage of KI pigs and WT pigs (Table 1 and Fig. 3D). These results were consistent with the one from randomly integrated PPARγ transgenic pigs. To further analyze intramuscular fat content, we performed H&E staining on soleus, gastrocnemius and longissimus dorsi of KI and WT pigs. The results showed that the connective and adipose tissues between the muscle fibers were also increased significantly in the muscles of KI pigs, compared to WT pigs (Fig. 3A). Moreover, the results of BODIPY and Oil Red O staining also showed the increase of IMF in longissimus dorsi of KI pigs (Fig. 3B and 3C).

**Table 1.** Comparison of carcass and meat quality traits between KI and WT pigs.

| Trait | KI (n=7) | WT (n=6) | *P* value |
| --- | --- | --- | --- |
| | Mean ± SD | Mean ± SD | |
| Slaughter age (day) | 159.42±3.96 | 158.67±3.77 | 0.3756 |
| Live weight (Kg) | 102.06±6.36 | 98.87±6.65 | 0.2170 |
| Carcass weight (Kg) | 72.54±4.16 | 71.30±7.83 | 0.3719 |
| Dressing percentage (%) | 71.13±1.93 | 72.02±4.79 | 0.3435 |
| Carcass length (cm) | 94.70±2.01 | 93.77±1.43 | 0.2011 |
| Average back fat thickness (mm) | 24.27±3.02 | 23.92±5.66 | 0.4486 |
| Average skin thickness (mm) | 3.01±0.21 | 3.08±0.38 | 0.3558 |
| Rib numbers | 15.00±0 | 15.00±0 | - |
| Loin eye area (cm<sup>2</sup>) | 38.34±2.13 | 37.86±3.17 | 0.3840 |
| Lean meat percentage (%) | 58.50±1.11 | 58.80±3.27 | 0.4188 |
| Fat meat percentage (%) | 25.37±1.89 | 24.40±3.20 | 0.2732 |
| Skin percentage (%) | 5.23±0.44 | 5.30±0.34 | 0.3849 |
| Bone percentage (%) | 10.91±1.09 | 11.50±0.32 | 0.1339 |
| Meat color score | 3.50±0.00 | 3.42±0.19 | 0.1498 |
| Meat marbling score | 2.86±0.35 | 2.42±0.19 | 0.0136* |
| L* | 44.52±0.72 | 46.64±1.34 | 0.0034** |
| a* | 7.14±1.25 | 7.95±1.99 | 0.2150 |
| b* | 11.42±0.4 | 12.38±1.57 | 0.0905 |
| Meat PH<sub>24</sub> | 5.88±0.20 | 5.80±0.18 | 0.2432 |
| Drip loss 48h (%) | 2.08±0.34 | 2.39±0.85 | 0.2138 |
| Water holding capacity (%) | 93.53±0.69 | 92.86±1.74 | 0.2026 |
| Water moisture (%) | 74.17±0.62 | 74.29±0.28 | 0.3567 |
| Intramuscular fat (%) | 2.82±0.28 | 2.35±0.16 | 0.0036** |

Results are presented as mean ± SD. \* $p$ <0.05 and \*\* $p$ <0.01 by non-paired Student's *t*-test.

**Fig. 3.**
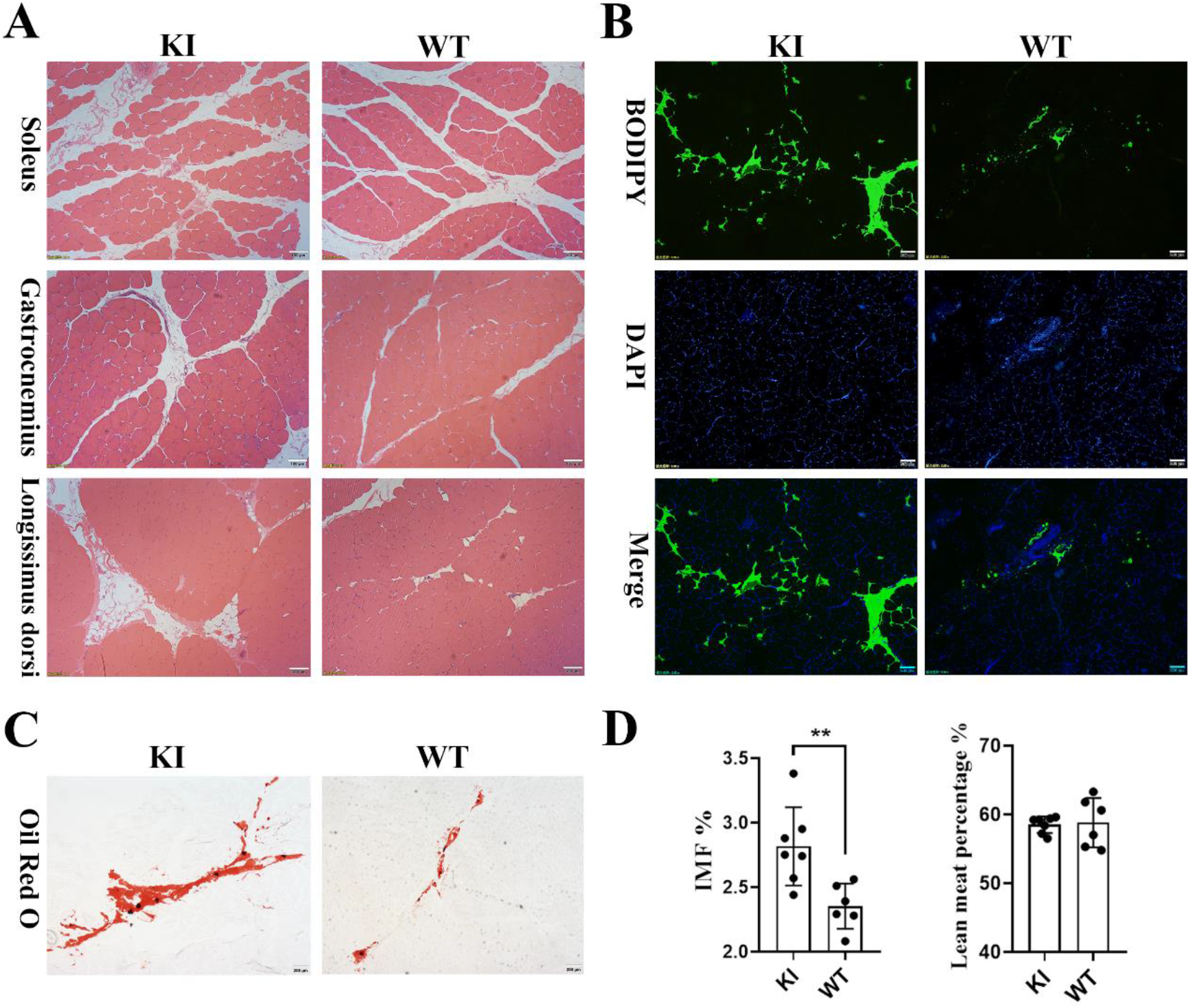
Muscle specific overexpression of PPARγ improves IMF in KI pigs. (A) Representative images of H&E staining for soleus muscle, gastrocnemius muscle and longissimus dorsi of KI and WT pigs. Scale bar: 100 μm. (B, C) Representative images of BODIPY (B) and Oil red O (C) staining for longissimus dorsi of KI and WT pigs. Scale bar: 200 μm. (D) Statistical results of IMF and lean meat percentage of KI (n=7) and WT (n=6) pigs. The data are presented as mean ± SD of independent experiments; * *p* < 0.05, ** *p* < 0.01. Dot plot reflects data points from independent experiment.

Given the critical role of PPARγ in adipogenesis, then we detected the downstream genes of PPARγ in the longissimus dorsi. The mRNA expression of LPL, CD36, FATP1, FABP4, PLIN1, and PLIN5 were all significantly increased in PPARγ KI pigs (Fig. 4A). Western blotting showed there was significant enhanced expression of LPL and FABP4 (Fig. 4B and 4C). We isolated the primary intramuscular preadipocytes from KI piglets and WT piglets, the results of Oil Red O staining showed more lipid droplets in PPARγ KI pigs (Fig. 4D). PPARγ, LPL and FABP4’s expression was also significantly upregulated in the primary intramuscular preadipocytes of KI pigs (Fig. 4E).

**Fig. 4.**
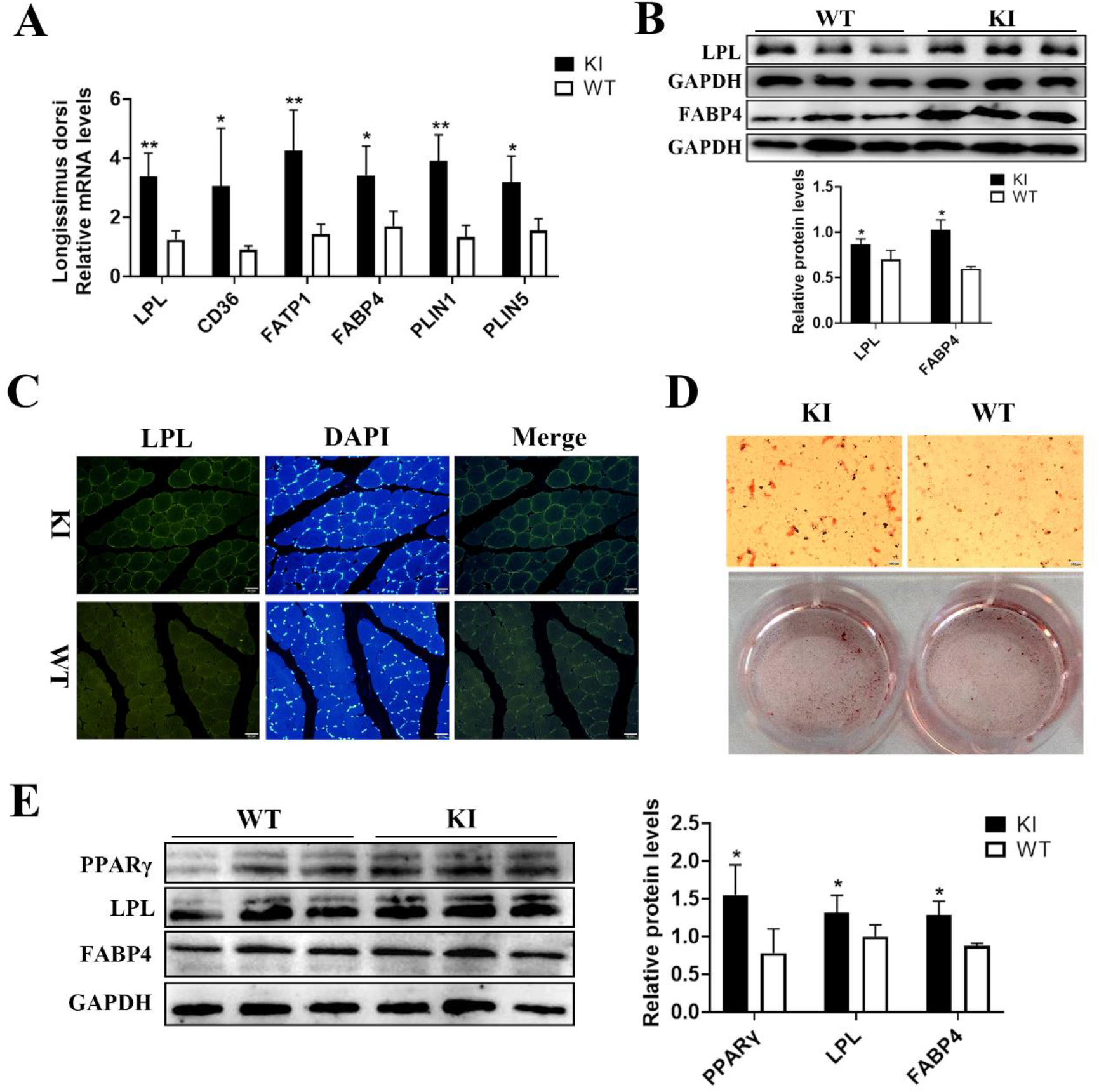
PPARγ induces the lipid metabolism genes’ expression in KI pigs. (A) Relative mRNA expression levels of genes induced by PPARγ in longissimus dorsi of KI and WT pigs. (B) Protein expression levels of LPL and FABP4 in longissimus dorsi of KI and WT pigs. (C) LPL immunofluorescence staining in Gastrocnemius muscles. Scale bar: 50 μm. (D) Oil red O staining of differentiated intramuscular preadipocytes isolated from KI and WT pigs. Scale bar: 100 μm. (E) Protein expression levels of PPARγ, LPL and FABP4 in differentiated intramuscular preadipocytes of KI and WT pigs (n=3). The relative mRNA and protein levels were normalized to GAPDH. The data are presented as mean ± SD of independent experiments; * *p* < 0.05, ** *p* < 0.01.

### 4. Muscle specific overexpression of PPARγ increased the proportion of oxidative muscle fibers

To further understand the influence of PPARγ specific overexpression in muscles, we performed label-free quantitative proteomics analyses in longissimus dorsi from 5 KI pigs and 5 WT pigs. A total of 108 differentially expressed proteins (DEPs) were found, of which 101 were up-regulated and 7 down-regulated (Fig. 5A). The KEGG predicted 107 enriched pathways, we showed the top 20 pathways in Fig. 5B. Most of DEPs were involved in the metabolic pathway, with a total of 24 up-regulated proteins (Fig. 5C). These proteins were involved in the process of fatty acid β-oxidation, citrate cycle (TCA cycle) and oxidative phosphorylation. They were key enzyme proteins or key constituent proteins in these processes, respectively (Fig. 5D). These up-regulated proteins in KI pigs were mainly involved in oxidative metabolism, especially the fatty acids oxidation. It is interesting that the marker genes of slow oxidative fiber MyHC1, TNNI1, and two constituent proteins of oxidative fibers (MYBPC1 and MYL2), were significantly increased in PPARγ KI pigs (Fig. 6A). Western blotting of MyHC1 and TNNI1 confirmed the proteomics results (Fig. 6C). ATPase staining of gastrocnemius showed that the proportion of slow fibers was significantly increased in KI pigs (Fig. 6D). In addition, real-time PCR analyses of three different muscles showed a consistently increasing expression of slow oxidative fiber related genes (Fig. S6 C). Further detection in the PPARγ TG pigs with random integration also confirmed this expression pattern (Fig. S6 B). Taken together, these results indicated that there were increased oxidative fibers in the muscle of PPARγ transgenic pigs.

**Fig. 5.**
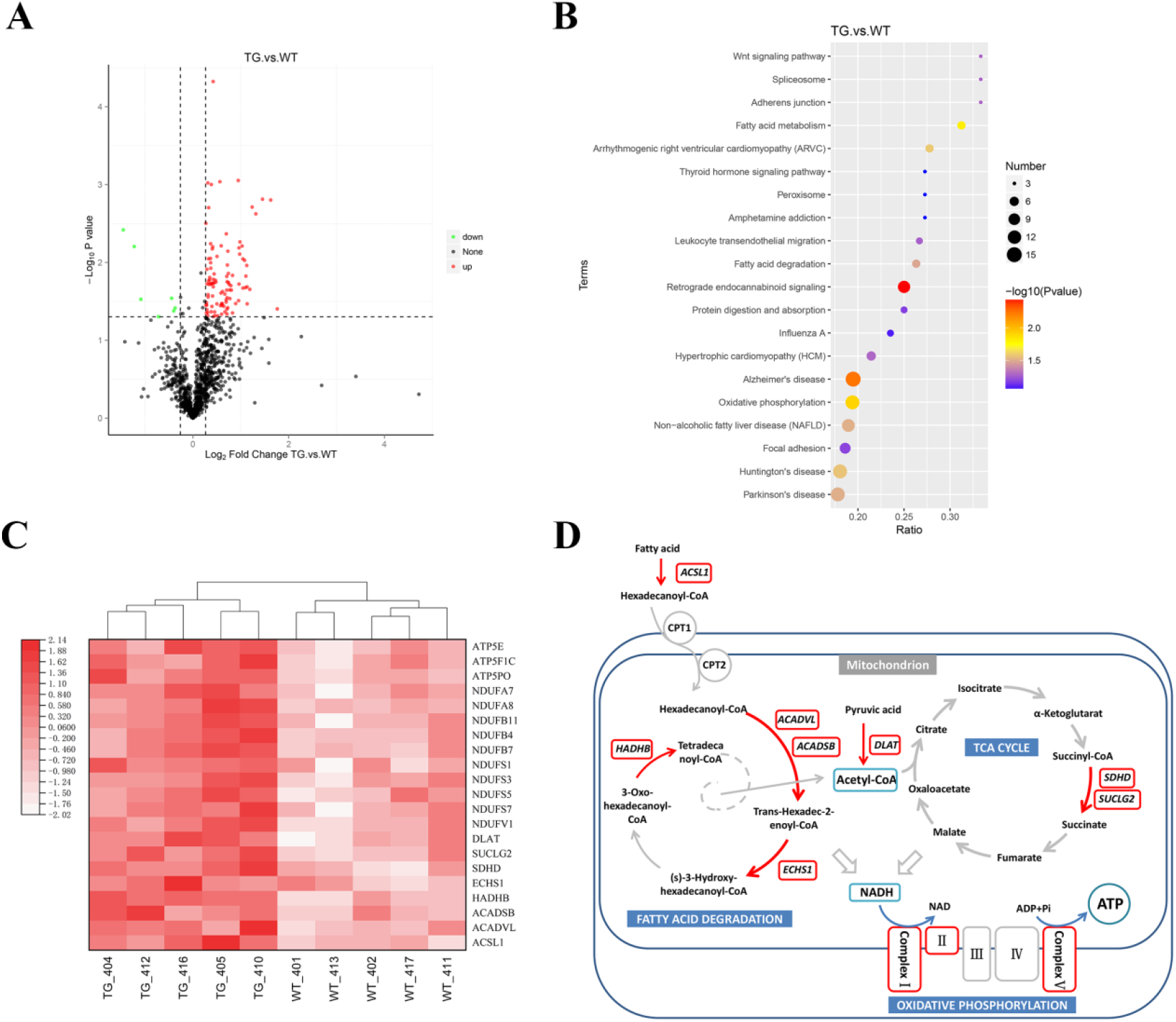
Proteomics analyses of longissimus dorsi muscles between KI pigs and WT pigs. (A) Volcano plot of differential expressed proteins. A total of 108 differentially expressed proteins were detected, of which 101 were up-regulated and 7 down-regulated. (B) Top 20 enriched pathways of differentially expressed proteins. The cycle size represents the number of proteins, and the color represents the P value. (C) The up-regulated proteins involved in the metabolic pathway, and (D) these proteins were involved in the process of fatty acid β-oxidation, citrate cycle (TCA cycle) and oxidative phosphorylation. Red boxes and red arrows represent key enzyme proteins or key constituent proteins in these processes.

**Fig. 6.**
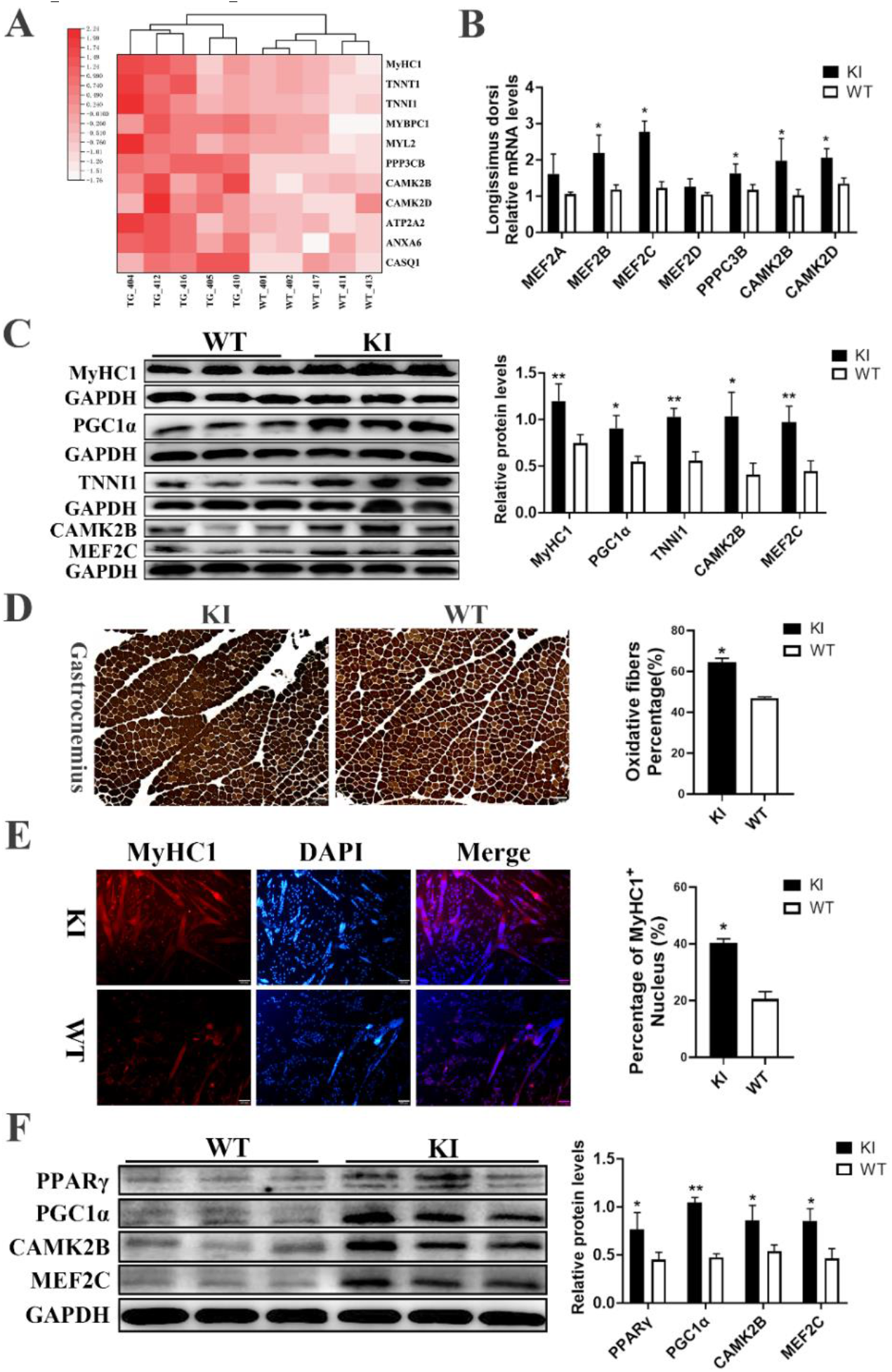
Overexpression of PPARγ in muscles increased the formation of slow oxidative muscle fibers. (A) The up-regulated proteins by proteomics which are involved in the calcium signaling pathway or related to oxidative muscle fiber. (B) Real-time PCR and (C) western blot method was used to test relative expression levels in KI and WT longissimus dorsi muscle. (D) ATPase staining in the Gastrocnemius muscle of KI and WT pigs. Scale bar: 200 μm. The dark color indicated slow oxidative fibers; the light color indicated glycolytic fibers. (E) MyHC1 immunofluorescence staining in differentiated satellite cells isolated from muscles of KI and WT pigs. Scale bar: 100 μm. (F) Western blot of differentiated satellite cells isolated from muscles of KI and WT pigs. The relative mRNA and protein levels were normalized to GAPDH. The data are presented as mean ± SD of independent experiments; * *p* < 0.05, ** *p* < 0.01.

### 5. Ca2+ mediated the oxidative muscle fiber switch in PPARγ transgenic pigs

To investigate the possible mechanisms in the formation of the oxidative muscle fibers regulated by PPARγ, we noticed that some typical calcium-dependent proteins are enriched in the calcium signaling pathway. In this pathway, PPP3CB (Protein Phosphatase 3 Catalytic Subunit Beta), CAMK2B (Calcium/Calmodulin Dependent Protein Kinase 2 Beta) and CAMK2D (Calcium/Calmodulin Dependent Protein Kinase 2 Delta) were all up-regulated in KI pigs (Fig. 6A), similar trend was observed by real-time PCR and western blot data (Fig. 6B and 6C). In our research, the expression levels of MEF2B, MEF2C and PGC1α were all significantly increased in muscles of KI pigs (Fig. 6B and 6C). The results of MyHC1 immunofluorescence and western blotting in satellite cells isolated from KI and WT pigs further confirmed that PPARγ overexpression promotes gene expression related to oxidative muscle fibers (Fig. 6E and 6F). Since CAMK and MEF2 were all regulated by the Ca^2+^ concentration, we propose that the increase of oxidative muscle fibers in KI pigs was caused by the regulation of calcium signaling pathway, Ca^2+^-sensitive phosphatases and kinases may played an important role in the switch of muscle fiber types mediated by PPARγ overexpression in skeletal muscle of the transgenic pigs (Fig. 7).

**Fig. 7.**
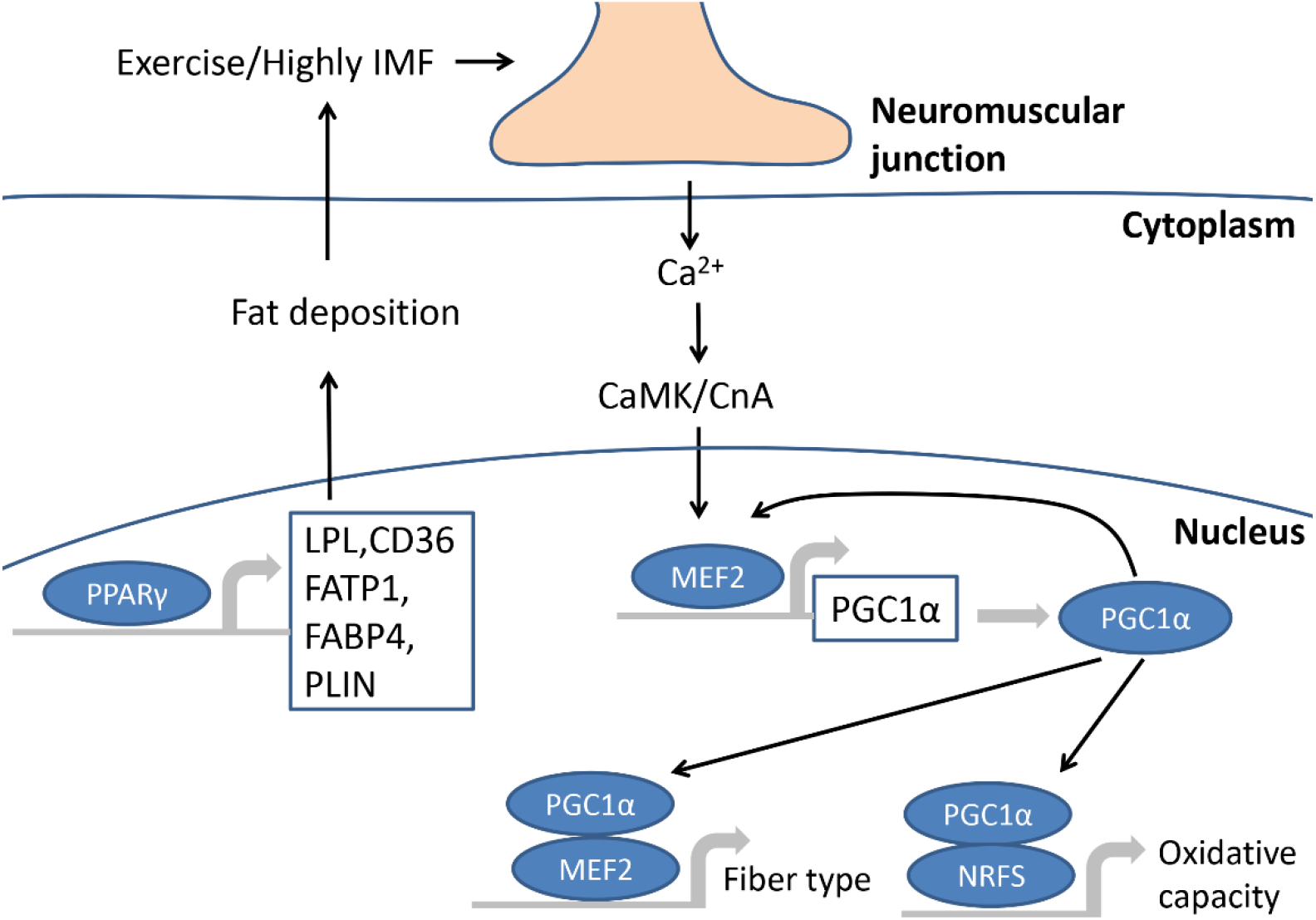
Schematic model of neuromuscular regulation mediated by endurance exercise or PPARγ overexpression in muscles. The overexpression of PPARγ can promote the fat deposition by induceing various genes. The metabolic requirements of exercise or highly IMF promotes the prolonged influx of Ca^2+^ by neuromuscular regulation. As calcium-dependent proteins, CAMK/CnA promotes the expression of MEF2 which can promote the expression of PGC1α. And as cofactor of MEF2, PGC1α further enhance the activity of MEF2. Through MEF2 and PGC1α, muscle specially overexpression of PPARγ can promotes the conversion of muscle fiber types and also participates in the promotion of oxidative capacity.

## Discussion

In recent years, there has been two types of methodologies to generate transgenic pigs by animal cloning procedures: random genomic insertion of the transgene, or specific targeted insertion of the transgene. As random insertion could land the transgenic construct anywhere in the genome, while CRISPR/Cas9 provides a precise and highly efficient method to integrate the transgene into a specific Rosa26 as safe harbor locus. As Rosa26 is a transcriptionally active locus that has been used over 700 times in the generation of transgenic animals based on Mouse Genome Informatics, it could make sure the transgene will be expressed, that site-specific integration will be influenced by random factors such as the silence of endogenous genes (Ruan et al., 2015). Randomly inserted transgenes may be unstable which can lead to changes in expression and changes in copy number, while the transgene will be there generation after generation by specific targeting into the safe harbor. Although the copy number of random integration pigs tended to be stable by hybridization with WT pigs. In our study, the KI pigs had one copy of insertion among generations, demonstrated the genetic stability of inserted genes. Additionally, neomycin gene in three of the six founder KI pigs had been successfully removed by adding adenovirus-Cre. The potential off-target effects (OTs) cannot be ignored in CRISPR/Cas9 mediate gene modified animals (Cho et al., 2014), while no off-target cleavages were identified for all 11 potential OT sites in our study. These data indicated that CRISPR/Cas9 specific targeted PPARγ pig model has good genetic stability and safety which without any labelled gene or uncontrollable randomness, such as unstable phenotypes, gene silencing or high copy number.

As a transcription factor, PPARγ activates multiple genes’ expression which involved in fat deposition, including LPL, FATP1, CD36, FABP4 and PLIN by binding a functional peroxisome proliferator response element (PPRE) on their promoter in adipose tissue (Dalen et al., 2004; Motojima et al., 1998; Schoonjans et al., 1996; Tontonoz et al., 1994). LPL is secreted by adipocytes and catalyzes the hydrolysis of triglycerides to free fatty acids (FFA) at capillary endothelium (Applebaum-Bowden, 1995; Kersten, 2014). Fatty acid transporters FATP1 and CD36 mediate the entry of FFA into adipocytes (Frohnert et al., 1999; Schaffer et al., 1994). FABP4 can bind to FFA with high affinity and act as lipid transport protein in adipocytes (Tontonoz et al., 1994). Abundant FFA in adipocytes will resynthesize triglycerides and stored in lipid droplets. PLIN is a protein family covering the surface of lipid droplets and keep the stability of lipid droplets by protecting them from lipases (Sztalryd et al., 2017). These genes’ expression was enhanced in PPARγ KI pigs (Fig.4) indicated the promotion effect of PPARγ on the accumulation of fat in myofibers and intramuscular adipocytes is similar to that in adipose tissue.

IMF is an important pork meat quality trait, which can influence tenderness, juiciness and the flavor of pork meat, but it is difficult to increase IMF and lean meat content at the same time by traditional breeding methods. Previous studies have shown that IMF is associated with PPARγ expression (Chen et al., 2011; Grindflek et al., 2001). Therefore, we investigated whether overexpression of PPARγ in muscles can promote the increase of IMF. In this study, we generated randomly integrated PPARγ TG pigs and site-specific integrated PPARγ KI pigs, respectively. Both of them can effectively achieve the overexpression of PPARγ in muscles, confirming both 7kb full-length swine MCK promoter and 1kb swine MCK promoter have the similar characteristics of muscle specific expression. Compared with wild-type pigs, PPARγ Transgenic pigs have the higher IMF while didn’t change the lean meat percentage or induce abnormalities in other tissues. As a commercial breed, White pig is well-known by its fast growth rate and high lean percentage, but the meat quality is not impressive. We used Large White pigs from two different sources for the propagation of PPARγ random transgenic and KI pigs, so their wild-type IMF had some difference, but the IMF was increased in both PPARγ transgenic pigs when compared with their own wild-type pig controls. And PPARγ pigs had higher meat marbling score. These results suggested that ectopic overexpression of PPARγ in muscles by transgenic technology is an effective approach to improve IMF and other meat quality traits.

Skeletal muscle fiber type can be switched when need to adapt to certain environmental cues. Slow fibers have more mitochondria, higher oxidative metabolizing enzyme activity and higher intracellular calcium concentration, are adapted for long and slow contraction (Holloszy et al., 1984; Westerblad et al., 1991). Muscle contraction is the activation of tension generation in muscle fibers. We elucidated a potential neuromuscular regulatory pathway that IMF can affect muscle fiber type conversion by proteomic sequencing. We found gene expression levels of PPP3CB, CAMK2, PGC1α and MEF2 were enhanced in PPARγ KI pigs. PPP3CB also known as calcineurin A Beta, is an important catalytic subunit of Calcineurin A (CnA). CAMK2B and CAMK2D are important subunits of CAMK2. CnA and CAMK2, as calcium-dependent proteins, play an important role in the process of increasing oxidative muscle fibers during endurance exercise. Studies have shown that the prolonged influx of Ca^2+^ into muscle cells during endurance exercise can effectively improve the activity of calmodulin-dependent kinases (CAMK2 and 5) and Calcineurin A, both of which can effectively promote the expression of myocyte enhancer factor 2 (MEF2) (Black et al., 1998; Potthoff et al., 2007). MEF2 can promote the expression of PGC1α, which plays a key role in the conversion of muscle fiber types. And PGC1α can act as a cofactor of MEF2, further amplifies activity of MEF2 (Fig.7). We found PPARγ plays an important role in the muscle fiber conversion from fast to slow twitch by increasing IMF. The relationship between muscle fiber composition and IMF content is complex but high correlation was observed. Recent study showed that a functional regulatory sequence variant in porcine MYH3 led to large amount of slow myofibers in the muscles and excess accumulation of IMF (Cho et al., 2019). Slow muscle fiber contains higher lipid than fast muscle fiber. In slow fibers, lipid droplets accounted for 0.5% of the fiber volume, while in fast fibers, accounted for less than 0.1% (Howald et al., 1985). As PPARγ pigs had highly IMF accumulation, higher concentration of FFA in muscle cells promotes oxidative metabolism of muscle cells with FA as substrate of mitochondrial oxidation process to generate ATP instead of glycolysis.

Insulin resistance in type 2 diabetes, impaired glucose tolerance, and metabolic syndrome is a well-established risk factor for cardiovascular disease. PPAR is the key regulator of the genes encoding proteins involved in insulin resistance. PPARγ agonists are commonly used in the treatment of type 2 diabetes (Doshi et al., 2009). Specific knockout PPARγ in skeletal muscle can cause severe insulin resistance, also have some adverse effects on adipose tissue and liver (Hevener et al., 2003). Owing to the critical role of PPARγ on insulin resistance, we were particularly interested in whether overexpression of PPARγ in muscle would affect the insulin sensitivity in KI pigs. The results showed there was no significant difference in glucose homeostasis between KI and WT pigs through glucose tolerance test (GTT) (Fig. S6 D). This may be due to the regulatory roles played by liver in insulin-mediated glucose metabolism (Gale, 2001), and the none significant overexpression of PPARγ in the liver of the KI pigs.

Pigs are genetically and physiologically closer to humans than small mammals. Furthermore, the existing genetic manipulation tools enable the generation of a variety of pig models to reflect human diseases (Holm et al., 2016; Prather et al., 2013). SCNT in combination with CRISPR-Cas9 allows for genetic modifications of the endogenous pig genes (Li et al., 2020; Yang et al., 2015; Zheng et al., 2017; Zhou et al., 2016). PPARγ agonists have also shown efficacy in Parkinson disease, Alzheimer disease, brain injury, and ALS. They act on microglial cells and inhibit the microglial cells activation (Medrano-Jiménez et al., 2019). PPARγ plays a well-established role in the regulation of lipid and glucose metabolism. It also has been demonstrated to modulate inflammation. For example, PPAR agonists inhibit the production of monocyte elaboration of inflammatory cytokines (Jiang et al., 1998). Additionally, PPAR receptor agonists have proven effective in suppressing the development of animal models of neuroinflammatory and neurodegenerative disorders (Zolezzi et al., 2017). The PPARγ agonist pioglitazone was demonstrated to modulate inflammation and induce neuroprotection in Parkinsonian monkeys, which supports the concept that PPARγ is a viable target against neurodegeneration (Swanson et al., 2011). Various neurodegenerative diseases are associated with electron transport chain enzyme activity reductions and increased mitochondrial-generated oxidative stress. Since our results showed 13 genes like NDUF families which involved in Parkinson disease, Alzheimer disease and Huntington’s disease were upregulated in our PPARγ2 KI pigs, implies our MCK-PPARγ2 transgenic pigs could be a large animal biomedical model for studying these human diseases.

In conclusion, skeletal muscule specific overexpression PPARγ promotes the formation of oxidative muscle fibers and increase the intramuscular fat content in pigs. In the skeletal muscle, Ca^2+^ mediated signaling phosphatases and kinases may play a key role as the second messenger of regulation of myofiber switching and fat deposition when PPARγ activity is elevated in skeletal muscle. Skeletal muscle-specific overexpression of PPARγ, generated by both transgenic pig methods, can promote oxidative fiber formation and intramuscular fat deposition in pigs. The research results provide a novel and effective approach for the improvement of meat quality traits in pig breeding.

## Materials and Methods

### Animals

All experiments involving animals were performed according to the guidelines of Good Laboratory Practice, and animal feeding and testing were conducted in accordance with the National Research Council Guide for the Care and Use of Laboratory Animals and approved by the Institutional Animal Care and Use Committee at Huazhong Agricultural University. Pigs were raised at the breeding swine quality supervision and testing center of Wuhan China. During experiment the pigs were allowed ad libitum to feed and water.

### Plasmids

To construct pN1-MCK-PPARγ vector for random integration, 7kb porcine MCK promoter was obtained according to our previous work (Ying et al., 2016). The pig PPARγ2 coding sequence was synthesized according to NCBI (NM_214379.1). To construct the donor vector for site-specific integration, 1kb MCK promoter was amplified from 7kb full-length MCK promoter, and the same pig PPARγ2 coding sequence was used. The left homology arm was 992bp and terminated at the 18th base of sgRNA. The right homology arm was 1556bp, starting at the 19th base of the sgRNA. The CRISPR/Cas9 construct was obtained as previously described (Cong et al., 2013). Targeting sequences of sgRNAs were designed using Cas-OFFinder online software (http://www.rgenome.net/cas-offinder/). The sgRNA sequences for knock-in were 5’-AGTTTGCTCCTTCTCGATTATGG-3’.

### Pig embryonic fibroblasts (PEFs) Culture and Transfection

PEFs were isolated from 25-day-old Large White pig fetuses and cultured in Dulbecco’s Modified Eagle’s Medium (DMEM) supplemented with 15% fetal bovine serum (FBS; Gibco, Grand Island, NY, USA). Plasmids were transfected into PEFs by electrical transfection in the condition of 170V/15ms (BIO-RAD, Gene Pulser Xcell). Selection culture medium supplemented with 400 μg/ml G418 for random integration cells and 400 μg/ml G418, 100 μM GCV for site-specific integration cells.

### Somatic cell nuclear transfer (SCNT)

Large White oocytes collected at a local abattoir were flushed out and matured in vitro, then enucleated the oocytes and reconstructed with the fetal fibroblast transfected with transgene cell lines. The reconstructed embryos were cultured at 39°C with 5% CO2 for 14-16h until embryo transfer. Embryos were surgically transferred into the oviduct of a surrogate the day after observed estrus. The pregnancy was detected by ultrasonography 21 days after embryo transfer. All transgenic piglets were natural delivered.

### Southern blot

Genomic DNA was isolated from pig ear punches using a DNA kit (Qiagen) according to the manufacturer’s protocol. Southern blot was performed according to the manufacturer’s protocol of DIG HIGH prime DNA labeling and detection starter kit 1 (Roche). 10 µg aliquot of genomic DNA was digested overnight with DraIII. The probe primers used for PCR amplification were: forward 5’-ATTGACCCAGAAAGCGATGC-3’, reverse 5’-GTGGACGCCATACTTTAGGA-3’.

### Isolation and culture of porcine primary cells

Primary intramuscular preadipocytes were isolated from longissimus dorsi of three-day new born piglets. The isolation and in vitro differentiation of primary intramuscular preadipocytes were performed as described previously (Wang et al., 2015). Primary myoblasts were isolated from the hind limb skeletal muscles of five-day piglets. Primary myoblasts were isolated and cultured according to the previously published method (Metzger et al., 2020).

### Histology staining

Immunofluorescence staining was performed according to the previously published method (Zhu et al., 2017). The antibodies included LPL (sc-373759; 1:200; Santa Cruz Biotechnology), MyHC1 (sc-53089; 1:200; Santa Cruz Biotechnology), Pax7 (1:100; Developmental Studies Hybridoma Bank, USA), and a secondary antibody (anti-mouse CY3 and anti-rabbit FITC; Beyotime Biotechnology, China). For hematoxylin and eosin (H&E) staining, muscles and tissues were immersed in 4% paraformaldehyde. H&E of muscle and tissues sections were carried out according to the reported method (Zhu et al., 2017). ATPase staining of gastrocnemius muscle was performed according to previous reported methods (Zhang et al., 2017). For Oil red and BODIPY staining of longissimus dorsi frozen sections were carried out according to the reported methods respectively (Huang et al., 2014; Son et al., 2018). The images were visualized using optical microscope (BX53; OLYMPUS).

### Real-time PCR Analysis

Total RNA was isolated with Trizol reagent (Invitrogen, Carlsbad, CA, USA) according to the manufacturer’s instructions. Real-time PCR was performed using the SYBR qPCR Mix (Toyobo, Japan) in a Bio-Rad CFX96 Real-Time PCR system with reaction volumes of 20 μL. The primers sequences listed in Table S5. The relative RNA expression levels were calculated using the Ct(2^-ΔΔCt^) method (Livak et al., 2001).

### Western Blotting

Tissues were frozen in liquid nitrogen immediately after dissection and stored at - 80°C until further use. Total proteins from tissue or cell were extracted using RIPA buffer with 1% (v/v) phenylmethylsulfonyl fluoride (PMSF) (Beyotime Biotechnology, China). The antibodies included PPARγ (ab45036; 1:1000; Abcam), LPL (sc-373759; 1:200; Santa Cruz Biotechnology), FABP4 (ab92501; 1:1000; Abcam), MyHC1 (sc-53089; 1:200; Santa Cruz Biotechnology), PGC1α (sc-518025; 1:200; Santa Cruz Biotechnology), TNNI1 (ab231720; 1:1000; Abcam), MEF2C (ab211493; 1:1000; Abcam), CAMKIIB (GB111070; 1:1000; Servicebio) Tubulin (ab7291; 1:1000; Abcam) GAPDH (sc-47724; 1:1000; Santa Cruz Biotechnology). The protein levels were normalized to GAPDH/Tubulin, and densitometric quantification of western blot bands were calculated by ImageJ software.

### Carcass and meat quality traits measurements

Pigs were slaughtered when weighed around 100kg and fasted for 24h before slaughtering. The carcass traits include live weight, carcass weight, dressing percentage, carcass length, average back fat thickness, average skin thickness, loin eye area, rib numbers, lean meat percentage, fat percentage, skin percentage and bone percentage. Meat quality traits were analyzed by meat color score, meat color lightness (L*), redness (a*), yellowness (b*) values, meat marbling score, meat pH24, drip loss 48h, water holding capacity, water moisture and intramuscular fat. All the traits were measured by Breeding Swine Quality Supervision and Testing Center, Ministry of Agriculture and Rural Affairs, P. R. China, according to National Profession Standards (No. NY/T 821-2004, NY/T 1180-2006, NY/T 825-2004).

### Postprandial glucose

Front cavity vein blood samples were collected for glucose test. All the 5-month-age animals used for testing were first proceeded 16 hrs fasting, then measured the glucose level at time point of 0, 2, 4 and 6 hours after feeding.

### Proteomics analysis

The longissimus dorsi of 5 KI and 5 WT pigs were collected and total protein was extracted according to previous reported methods (Silva et al., 2019). Proteomics analyses were performed using an EASY-nLCTM 1200 UHPLC system (Thermo Fisher) coupled with an Orbitrap Q Exactive HF-X mass spectrometer (Thermo Fisher) operating in the data-dependent acquisition (DDA) mode. Protein identification was carried out according to the reported method (Silva et al., 2019). P≤0.05 and the absolute value of Ratio≥1.2 were set as the threshold to determine significant differentially expressed proteins (DEP). Pathway analysis was used to identify the significant pathways of the DEP according to KEGG database (http://www.genome.ad.jp/kegg/).

### Statistical Analysis

All data were expressed as mean ± SD. Differences between groups were analyzed by two-tailed Student’s t-test. P-values less than 0.05 were considered to represent statistically significant differences.

## Acknowledgments

We thank Hongxing Chen, Youliang Wang, Xiaojie Wu and Yanli Lin at Beijing Institute of Biotechnology for their technical supports in vector construction and sgRNA screening. We also thank Zhen Liu, Jinlong Zhang, Changwei Qiu, Guanwei Nie, and Jun Liu for their technical supports, breeding and raising the transgenic pigs. This work was financially supported by the National Key Project of Transgenic Research (Grant No. 2016ZX08006-002), the Agricultural Innovation Fund of Hubei Province (2016-620-000-001-043), the Fundamental Research Funds for the Central Universities (Program No. 2662018PY045).

## Author contributions

B.Z., J.Y., Y.Z.X., Y.Z., H.G. conceived and designed the research; Y.Z., H.G., G.B. performed experiments; J.L, Y.P, X.Z., Y.M., W.J., L.H., L.Z., T.L., X.X., Z.M., X.W., Y.L. helped with the generation of transgenic pigs. Y.Z., H.G. analyzed the data; Y.Z., H.G., B.Z., J.Y. wrote the manuscript. All authors read and approved the final manuscript.

