## Supplemental information for "Targeted over-expression of PPARγ in pig skeletal muscle improves oxidative fiber formation and intramuscular fat deposition"

**Fig. S1. Construction of PPARγ transgenic plasmid and identification of TG pigs.** (A) Schematic illustration of the transgenic plasmid which contains an expression cassette consisting of porcine PPARγ cDNA driven by a 7kb muscle tissue-specific MCK promoter and positive selection cassette consisting of a neomycin resistance gene (NeoR) driven by the SV40 promoter. The opposite red arrows are primers for PCR verification of TG pigs. (B) PCR identification of F1 generation TG pigs. (C) Southern blotting identification of F1 TG pigs. Lane N represents negative control; lane P represents positive control. (D) PCR identification of F2 generation TG pigs.

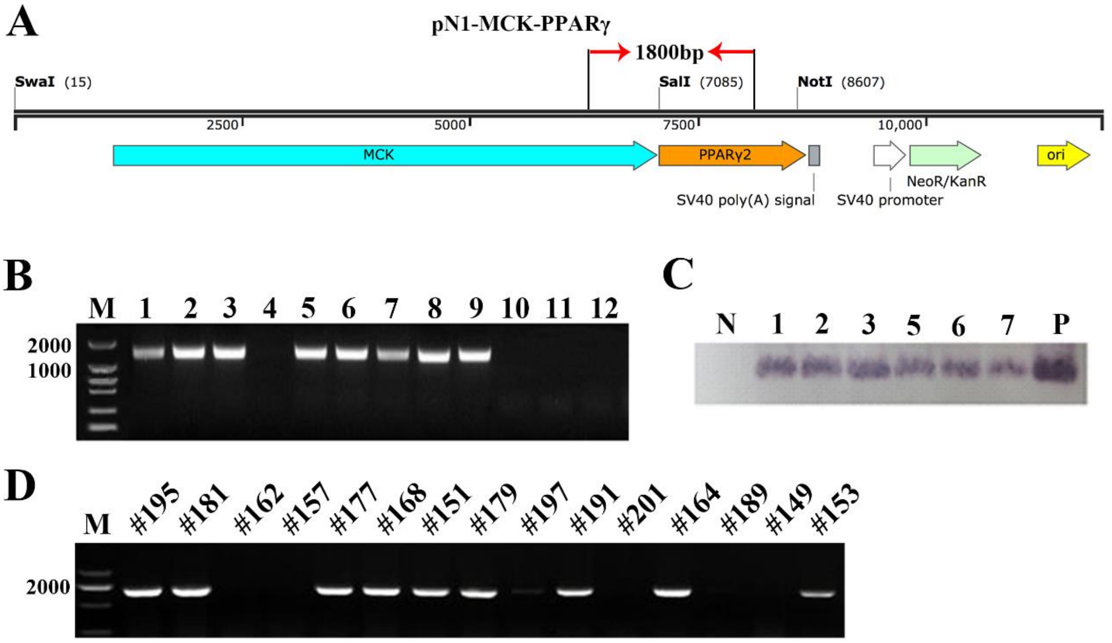

**Fig. S2. PCR identification and copy number detection of KI pigs.** (A) Sanger sequencing of the 5′junction and the 3′junction confirms the targeted insertion of exogenous DNA. (B) The site-specific PCR analyses of F1 generation KI pigs, which obtained by mating #314 founder boar with WT sow. Lanes #401-#417 represent 12 F1 pigs, of which #404, #405, #410, #412, #414 and #416 are KI pigs. Lane N represents negative control WT pigs. (C) The standard regression curve was drawn with the ΔCT of 8 standards containing 1, 2, 6, 8, 10, 16, 32 and 48 copies as the vertical axis and $\log_{10}$(copy number) as the horizontal axis ($R^2$=0.9939). And the copy numbers are calculated by this curve. Dot plot reflects data points from independent experiment.

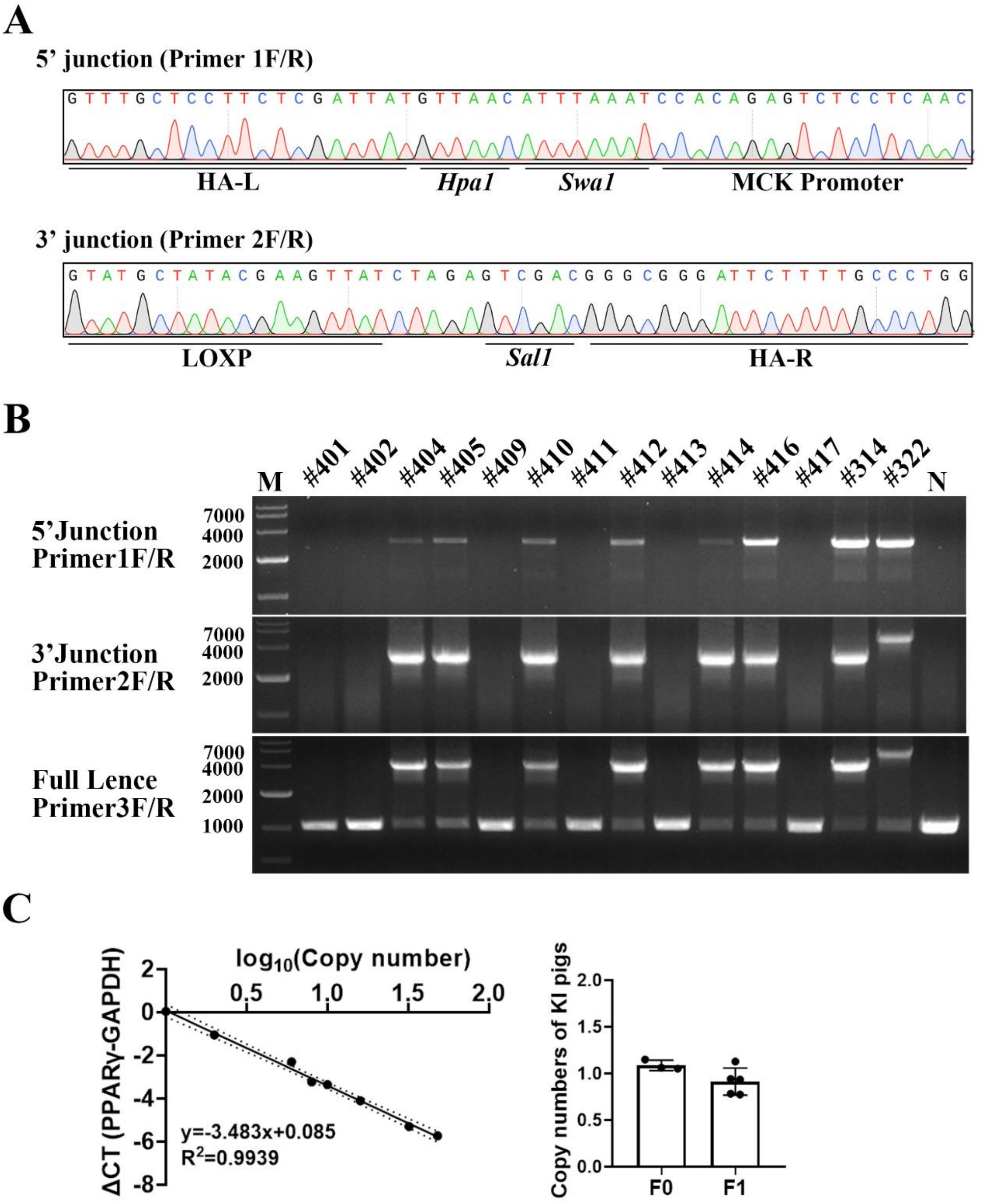

**Fig. S3. CRISPR/Cas9 mediated PPARγ KI has no off-target effects (OTs).** (A) Potential OT sites predicted by Cas-OFFinder online software. The green and red highlights represent the bases that match the sgRNA guidance sequence and PAM sequence, respectively. (B) PCR analyses of these 11 potential OT sites. 'a' for KI pigs and 'b' for WT pigs. The primers were listed in Table S5.

**A**

| | Location | 1 | 2 | 3 | 4 | 5 | 6 | 7 | 8 | 9 | 10 | 11 | 12 | 13 | 14 | 15 | 16 | 17 | 18 | 19 | 20 | P | A | M |
|---|---|---|---|---|---|---|---|---|---|---|---|---|---|---|---|---|---|---|---|---|---|---|---|---|
| SgRNA | CM000824.5-65758620 | G | C | T | C | C | T | T | C | T | C | G | A | T | T | A | T | G | G | G | C | G | G | G |
| 1 | CM000817.5-38168686 | G | C | T | C | C | T | C | C | T | C | C | A | T | T | C | T | G | G | G | C | T | G | G |
| 2 | CM000818.5-7460631 | G | C | T | C | C | T | T | C | T | C | T | A | T | G | T | T | G | G | G | C | A | G | G |
| 3 | CM000822.5-78847602 | G | C | T | C | C | C | T | C | T | C | A | A | T | T | C | T | G | G | G | C | A | G | G |
| 4 | CM000825.5-60351861 | G | C | T | C | C | T | T | C | T | C | C | A | T | G | T | T | G | G | G | C | A | G | G |
| 5 | CM000829.5-34982114 | G | C | T | C | C | T | T | C | A | G | G | A | T | G | A | T | G | G | G | C | T | G | G |
| 6 | CM000812.5-119006866 | G | G | T | C | C | T | G | C | T | C | A | G | T | T | A | T | G | G | G | C | A | G | G |
| 7 | CM000814.5-124626688 | G | G | T | C | C | T | T | G | T | C | T | T | T | T | A | T | G | G | G | C | C | G | G |
| 8 | CM000815.5-130352333 | G | C | T | C | A | T | G | C | G | A | G | A | T | T | A | T | G | G | G | C | T | G | G |
| 9 | CM000820.5-52539316 | G | C | C | C | T | T | T | C | C | C | A | A | T | T | A | T | G | G | G | C | A | G | G |
| 10 | CM000822.5-22802563 | C | C | T | C | C | T | G | C | T | C | T | T | T | T | A | T | G | G | G | C | C | G | G |
| 11 | CM000815.5-6973644 | G | C | T | C | T | A | T | C | C | T | G | A | T | T | A | T | G | G | G | C | T | G | G |

**B**

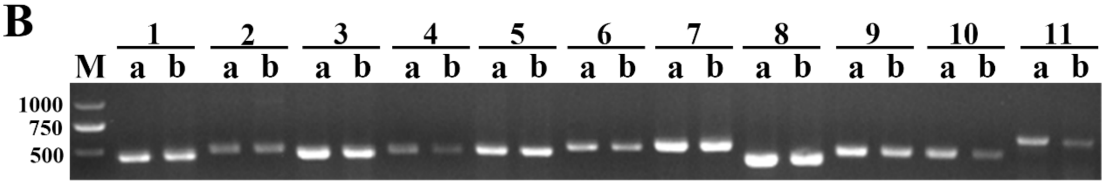

**Fig. S4. Muscle specific overexpression of PPARγ in KI pigs.** (A) The relative PPARγ mRNA expression levels in different tissues of F0 KI pigs compared with WT pigs (n=3). (B) Western blotting results showed there was no significant difference for PPARγ protein levels in heart and liver of KI and WT pigs (n=3). The relative mRNA and protein levels were normalized to GAPDH. The data are presented as mean ± SD of independent experiments; * $p < 0.05$, ** $p < 0.01$.

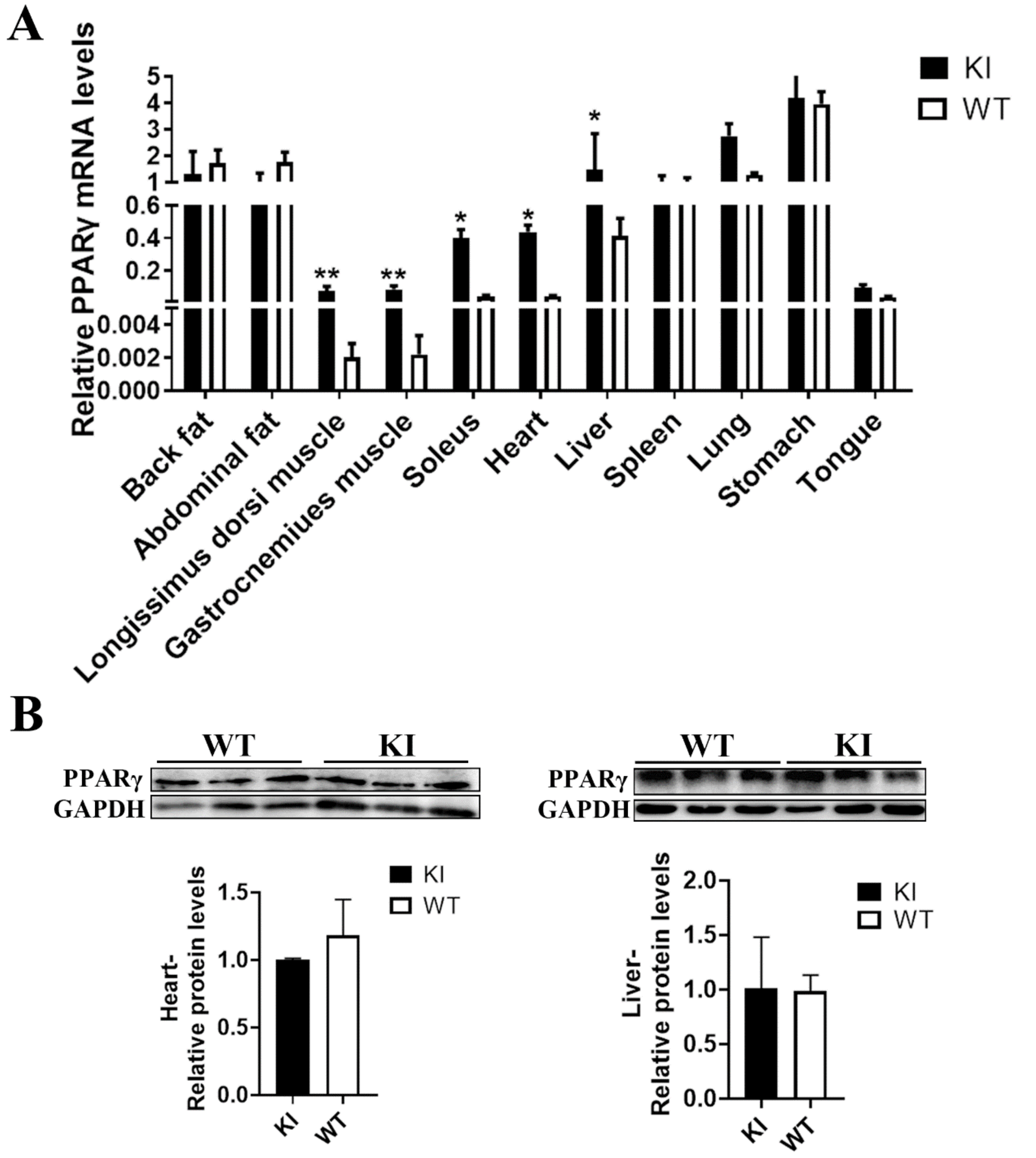

**Fig. S5. Muscle overexpression of PPARγ in KI pigs does not affect tissue morphology and weights.** (A) H&E staining of heart, liver, spleen, lung, kidney and backfat showed there was no significant difference for morphology of these tissues between KI and WT pigs. Scale bar: 100 μm. (B) There was no significant difference for weight of tissues between KI and WT pigs (n=5).

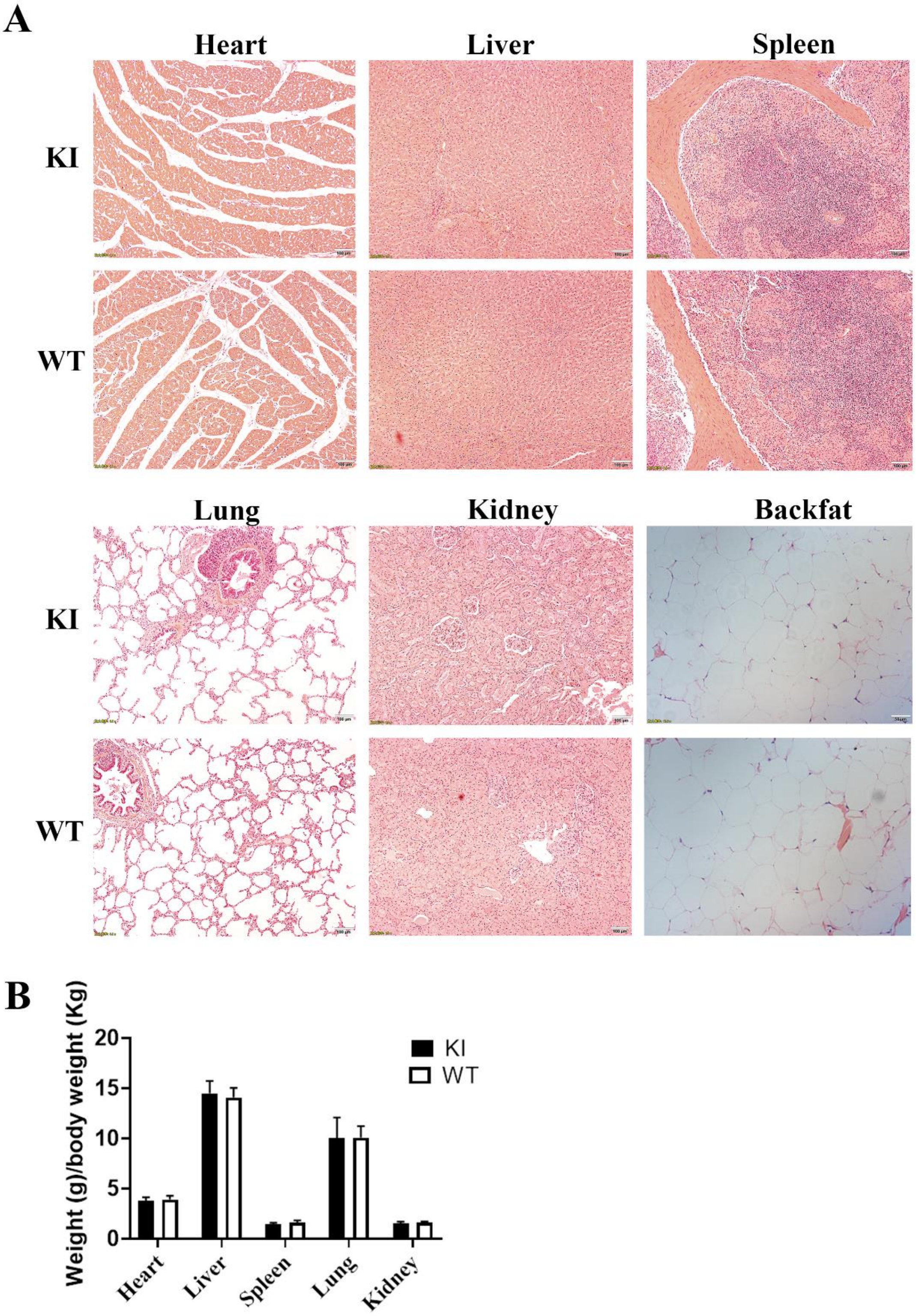


**Fig. S6. Muscle overexpression of PPARγ increased gene expression related to**

**oxidative muscle fibers.** (A) Growth curve showed there was no difference for body weights between KI (n = 7) and WT (n = 6) pigs. (B) Relative mRNA expression levels in soleus muscle of TG pigs (n=3). (C) Relative mRNA expression levels in longissimus dorsi, gastrocnemius and soleus. (D) Glucose concentration curve of the plasma after 0h, 2h, 4h and 6h after eating of KI and WT pigs (n=3). The relative mRNA levels were normalized to GAPDH. The data are presented as mean ± SD of independent experiments; * $p < 0.05$, ** $p < 0.01$.

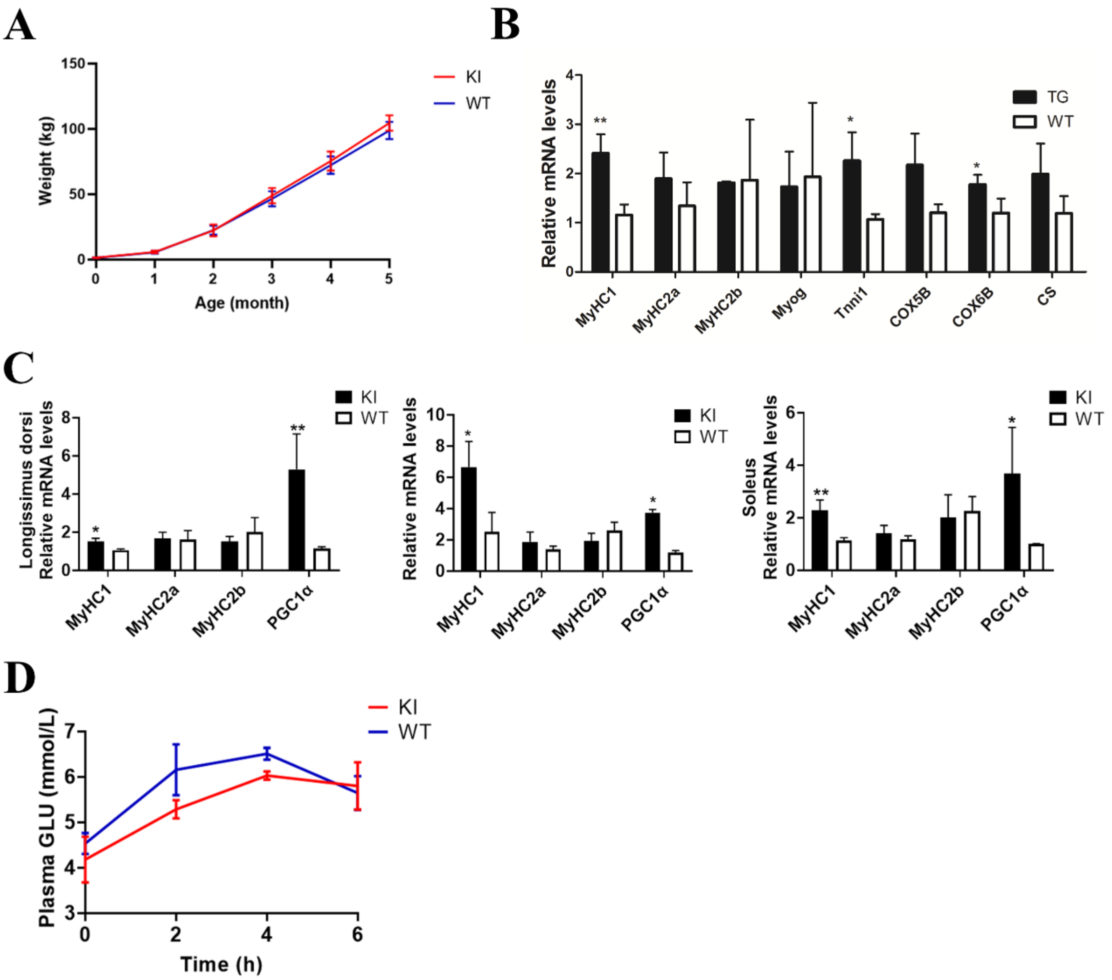

Table S1 Comparison of carcass and meat quality traits between F1 TG and WT pigs

| Traits | TG pigs (n=7) (Mean ± SD) | WT pigs (n=7) (Mean ± SD) | *P* values |
|---|---|---|---|
| Slaughter age (day) | 194.9±4.5 | 196.1±4.5 | 0.313 |
| Live weight (Kg) | 116.0±4.8 | 110.2±11.9 | 0.231 |
| Carcass weight (Kg) | 88.2±5.0 | 83.2±10.4 | 0.457 |
| Dressing percentage (%) | 75.9±1.2 | 75.2±1.5 | 0.285 |
| Carcass length (cm) | 97.0±1.4 | 97.5±3.1 | 0.156 |
| Average back fat thickness (mm) | 23.9±0.6 | 18.8±2.2 | 0.278 |
| Average skin thickness (mm) | 4.4±0.1 | 3.8±0.4 | 0.375 |
| Rib numbers (pair) | 15.0±0.0 | 15.3±0.3 | 0.169 |
| Loin eye area ($cm^2$) | 49.4±3.0 | 54.1±7.0 | 0.185 |
| Lean meat percentage (%) | 67.4±1.0 | 67.8±1.2 | 0.178 |
| Fat percentage (%) | 14.1±1.1 | 12.3±2.1 | 0.413 |
| Skin percentage (%) | 7.8±0.2 | 8.1±0.4 | 0.128 |
| Bone percentage (%) | 10.7±0.5 | 11.8±0.7 | 0.357 |
| Meat marbling score | 2.3±0.2 | 1.7±0.4 | 0.068 |
| L* | 40.1±2.9 | 41.1±2.5 | 0.175 |
| a* | 8.3±2.0 | 9.8±1.4 | 0.089 |
| b* | 13.6±0.5 | 14.7±0.9 | 0.134 |
| Meat $pH_{24}$ | 5.7±0.1 | 5.7±0.0 | 0.091 |
| Drip loss 48h (%) | 1.9±0.4 | 2.9±0.4 | 0.107 |
| Water holding capacity (%) | 94.2±0.4 | 94.2±0.2 | 0.256 |
| Water moisture (%) | 73.7±0.2 | 73.7±0.2 | 0.378 |
| Intramuscular fat (%) | 2.3±0.2 | 1.5±0.2 | 0.016* |

Table S2 Comparison of carcass and meat quality traits between F2 TG and WT pigs

| Traits | TG pigs (n=6) (Mean ± SD) | WT pigs (n=6) (Mean ± SD) | *P* values |
|---|---|---|---|
| Slaughter age (day) | 164.3±0.94 | 163.7±0.94 | 0.145 |
| Live weight (Kg) | 99.8±6.0 | 97.7±7.7 | 0.381 |
| Carcass weight (Kg) | 74.5±5.0 | 71.7±5.7 | 0.218 |
| Dressing percentage (%) | 74.6±1.1 | 73.4±2.3 | 0.197 |
| Carcass length (cm) | 93.7±1.5 | 93.9±2.9 | 0.211 |
| Average back fat thickness (mm) | 21.4±5.4 | 15.2±5.8 | 0.138 |
| Average skin thickness (mm) | 4.0±0.9 | 3.9±1.0 | 0.233 |
| Rib numbers | 15.0±0.0 | 15.0±0.0 | — |
| Loin eye area ($cm^2$) | 42.6±6.5 | 45.7±6.4 | 0.343 |
| Lean meat percentage (%) | 65.2±3.3 | 67.7±3.4 | 0.258 |
| Fat percentage (%) | 16.7±3.5 | 13.0±2.7 | 0.345 |
| Skin percentage (%) | 6.8±0.9 | 7.2±1.1 | 0.256 |
| Bone percentage (%) | 11.1±0.6 | 12.1±0.6 | 0.117 |
| Meat color score | 3.3±0.3 | 3.2±0.3 | 0.458 |
| Meat marbling score | 1.6±0.6 | 1.2±0.2 | 0.067 |

| | | | |
|---|---|---|---|
| Meat $pH_{24}$ | 5.6±0.1 | 5.6±0.1 | 0.258 |
| Drip loss 48h (%) | 2.2±0.4 | 2.2±0.7 | 0.288 |
| Water holding capacity (%) | 92.7±0.5 | 92.4±1.1 | 0.312 |
| Water moisture (%) | 74.6±0.5 | 74.6±0.8 | 0.122 |
| Intramuscular fat (%) | 2.4±0.3 | 1.7±0.1 | 0.008* |

Table S3. Summary of embryo transfer results of KI pigs

| | Pig ID of surrogate | Embryos transferred | Pregnancy | Piglets at birth | KI piglets | Survived for 1 month | Neomycin free pigs |
|---|---|---|---|---|---|---|---|
| | 46 | 345 | No | - | - | - | - |
| | 112 | 153 | No | - | - | - | - |
| | 124 | 246 | Yes | - | - | - | - |
| | 74 | 148 | No | - | - | - | - |
| | 142 | 168 | Yes | 4 | 4 | 1 | 1 |
| | 42 | 254 | Yes | 2 | 2 | 2 | 1 |
| | 122 | 157 | Yes | 3 | 3 | 0 | - |
| | 100 | 164 | Yes | 4 | 4 | 3 | 1 |
| | 112 | 176 | No | - | - | - | - |
| | 44 | 264 | No | - | - | - | - |
| Total | 10 | 2075 | 5 | 13 | 13 | 6 | 3 |

Table S4. Summary of F1 KI pigs

| Male parent of F0 | WT female parent | Piglets at birth | Male/female | KI/WT |
|---|---|---|---|---|
| #314 | #1 | 20 | 11/9 | 10/10 |
| | #4 | 12 | 7/5 | 6/6 |
| Total | 2 | 32 | 1.3:1 | 1:1 |

Table S5. Primers used for PCR

| Gene or Primer name | Primer sequence (5'-3') |
|---|---|
| 5' junction primer 1 | F: TATCGTTTGTTACGCTGGAAGGGGAAGA |
| | R: GGGAGTTTATTTTTAGAGCTCGCTACTCG |
| 3' junction primer 2 | F: CGCTCCGTGGAGGCCGTGCAGGAGATC |
| | R: GGTACAAGACTCAACAAGAACCTGTGCC |
| Full length primer 3 | F: TGATTGGCTGCTGAAGTCCTGGGAACGG |
| | R: GCCAATGCTATGTCTGGGACTGGATGAG |
| PPARγ | F: TCCCGCTGACCAAAGCAAAGGC |
| | R: CCACGGAGCGAAACTGACACCC |
| LPL | F: CAAACTTGTGGCTGCCCTAT |
| | R: GTGGACATTGTTGGGAGGAT |
| CD36 | F: ATCGTGCCTATCCTCTGG |
| | R: CCAGGCCAAGGAGGTTAA |
| FATP1 | F: ATCAACACCAACCTGCGG |

| | |
|---|---|
| | R: GAAGAGGCTGAGCGAGGG |
| FABP4 | F: CTGAGATTGCCTTCAAATTG |
| | R: CTTGGCTTATGCTCTCTCATA |
| PLIN1 | F: GCCTGACTTTGCTGGATGG |
| | R: CTTGGTGCTGGTGTAGGTCTTCT |
| PLIN5 | F: GTCTCCGATGCTTATAGCGC |
| | R: TGAAGGCTGCTGAAGAAAGG |
| MyHC1 | F: GAGGAAGAGTGAGCGTCGCAT |
| | R: ACCTTCAGCTGTAGCTTGTCCA |
| MyHC2a | F: GATGGAGATCGACGACCTTGCT |
| | R: CTGCTGCTCTTCCTCCTTGGAT |
| MyHC2b | F: CGCCAAGCTACTGAGGCAATAA |
| | R: GTTCCACCATGGCCAGTTGTTC |
| PGC1α | F: CGCAAGCTTCTCTGAGCTTCTTT |
| | R:GGATACACTTTGCGCAGGTCGAA |
| TNNI1 | F: GCTCTAAACACAAGGTGTCCAT |
| | R: GCCTCGACGTTCTTTCTCCAGT |
| Myoglobin | F: AGCACCTGAAGTCAGAGGATGA |
| | R: TCCAGGTACTTGACAGGGATCT |
| COX5B | F: CTATGGCATCTGGAGGTGGTGT |
| | R: CTATCCGCTTGTTGGTGATGGA |
| COX6B | F: GTCACATTGAGCTTCCAGCGGT |
| | R: AGCAGTCATTGCTTTCTCACAGC |
| CS | F: GGAAGTGCTTGTTTGGCTGACA |
| | R: CATGAGGCAGGTGTTTCAGAGCA |
| MEF2A | F: TGAATACCCAGAGGATAAGCAGTT |
| | R: TAATCGGTGTTGTAGGCGG |
| MEF2B | F: ACAATGGGGAGGAAAAAAAT |
| | R: CTGTTGAAGATGATGAGGGC |
| MEF2C | F: CAGGTGGTCTGATGGGTG |
| | R: CTGGTGGAATAAGAACTCG |
| MEF2D | F: GAACCGACAGGTGACATT |
| | R: TGGAGTGGTTGAAGATGA |
| PPPC3B | F: TAGTAAAAGAAGGTCGGGT |
| | R: ATCGTGTATTAGCAGGTGA |
| CAMK2B | F: GACCTCAAGCCCGAGAACCT |
| | R: TGCCAGCGAACCCAAACCAT |
| CAMK2D | F: GGGACTTGAAGCCTGAGAAT |
| | R: CTGCTGGTCTCCTTGAACTT |
| GAPDH | F: CCACAACATACGTAGCACCACGATC |
| | R: CCTTCATTGACCTCCACTACATGGT |
| OTs-1 | F: ATTCAGCCAGAGGCAAAGTG |
| | R: ATGCCCCCAAACAATAAAAC |
| OTs-2 | F: CCCTGAGAGACAGAGTGACA |

| | |
|---|---|
| | R: CAACAAATCCACACAACAGA |
| OTs-3 | F: CGTCTCACCTTCCTGGCATT |
| | R: CCTGGGGACCTGAGAACCTT |
| OTs-4 | F: GTAGATTTTTCTGTCTCGCA |
| | R: AAACAATAAAACAATGGTGG |
| OTs-5 | F: GAGGATAGTAGGGTCTGTGT |
| | R: GCAACTGTTCTCTTCTTTCT |
| OTs-6 | F: ATTGGTTGATGATGAGGTAACAAGGTGG |
| | R: CTCTCTCCAACTTGTCTTCGGCTGTCTC |
| OTs-7 | F: AGGTCCACCCCAACCACGCCCTCAAA |
| | R: TCACACTGTAGACACACTCCTGCCGA |
| OTs-8 | F: CTCTTGCTGTGCCTTTCCGA |
| | R: TGCCTGACCCCCTCTGAAT |
| OTs-9 | F: CTGCCCAGATTCTCCGATTG |
| | R: TCTTCCTATTCTCCCCTTCA |
| OTs-10 | F: CACTCTTACGACCCCATCTA |
| | R: ACTCTGGCTGTGAAATAAAG |
| OTs-11 | F: ATCCTAAAAGTTCTCATCAC |
| | R: AACTATGGTTACCGTTTACT |

OTs represents potential off-target sites.

### Date S1. Sequence of 3870bp full length PCR in #314 KI pig

*Start* (0)

CTACTACCCGCTTGGGTCCAATGAGCGAACCACTACGCGAGTCGGCAGGGAGGCGGTCTTTGGTACGGCCCTCCCCGAGGCCAGCGCCGC 90
GATGATGGGCGAACCCAGGTTACTCGCTTGGTGATGCGCTCAGCCGTCCCTCCGCCAGAAACCATGCCGGGAGGGGCTCCGGTCGCGGCG
HA-L

AGTGTCTGGCCCCTCGCCCCTGCGCAACGTGGCAGGAAGCGCGCGCAGGAGGCGGGGGCGGGCTGCCGGGCCGAGGCTTCTGGGTGGTGG 180
TCACAGACCGGGGAGCGGGGACGCGTTGCACCGTCCTTCGCGCGCGTCCTCCGCCCCCGCCCGACGGCCCGGCTCCGAAGACCCACCACC
HA-L

TGACTGCGGCTCCGCCCTGGGCGTCCGCCGCCTGAAGGACGAGACTAGCTCTCTACCTGCTCTCGGACCCGTGGGGGTGGGGGGTGGAGG 270
ACTGACGCCGAGGCGGGACCCGCAGGCGGCGGACTTCCTGCTCTGATCGAGAGATGGACGAGAGCCTGGGCACCCCCACCCCCCACCTCC
HA-L

AAGGAGTGGGGGGTCGGTCCCTGCTGGCTTGTGGGTGGGAGGCGCATGTTTCTCCAAAACCCGCGCGAGCTGCAATCCTGAGGGAGCTGC 360
TTCCTCACCCCCCAGCCAGGGACGACCGAACACCCACCCTCCGCGTACAAAGAGGTTTTGGGCGCGCTCGACGTTAGGACTCCCTCGACG
HA-L

AGTGGAGGAGGCGGAGAGAAGGCCGCACCCTTCTCCGCAGGGGGAGGGGAGTGCCGCAATACCTTTATGGGAGTTCTCTGCTGCCTCCTT 450
TCACCTCCTCCGCCTCTCTTCCGGCGTGGGAAGAGGCGTCCCCCTCCCCTCACGGCGTTATGGAAATACCCTCAAGAGACGACGGAGGAA
HA-L

TTCCTAAGGACCGCCCTGGGCCTAGAAAAATCCCTCCCTCCCCCGCGATCTCGTCATCGCCTCCATGTCAGTTTGCTCCTTCTCGATTAT 540
AAGGATTCCTGGCGGGACCCGGATCTTTTTAGGGAGGGAGGGGGCGCTAGAGCAGTAGCGGAGGTACAGTCAAACGAGGAAGAGCTAATA
HA-L

**SwaI**

GTTAACATTTAAATCCACAGAGTCTCCTCAACCCCGAGAGCCCTGTGCTCTGACCCGAGTTGCCGCCCAGCCTCCTCCCCCTGCGGGTGG 630
CAATTGTAAATTTAGGTGTCTCAGAGGAGTTGGGGCTCTCGGGACACGAGACTGGGCTCAACGGCGGGTCGGAGGAGGGGGACGCCCACC
MCK promoter
HA-L

GTGTGGACGCCTCCCCAGGGCCGGGGCTGTGGCTGCCCTTGTAAGGAGGTGAGGCCTGGGGACACCAGAGAGGCCTGGTTATAATTAACC 720
CACACCTGCGGAGGGGTCCCGGCCCCGACACCGACGGGAACATTCCTCCACTCCGGACCCCTGTGGTCTCTCCGGACCAATATTAATTGG
MCK promoter

GGGACACGTGGCCAGCCCGCCCCCAACACCTGCCCCCGCCCCTGCCCCCATCCCCAGCGCCTCGGGTCTCCCGGAGGAGACAGCGAGTAG 810
CCCTGTGCACCGGTCGGGCGGGGGTTGTGGACGGGGGCGGGGACGGGGGTAGGGGTCGCGGAGCCCAGAGGGCCTCCTCTGTCGCTCATC
MCK promoter

CGAGCTCTAAAAATAAACTCCCTTTTCTGCAAGCCTGCAGGCCCTGTCCCCTCCAGCGTGGAATCACCCAGTGTCACTGGGCCCTGCGCC 900
GCTCGAGATTTTTATTTGAGGGAAAAGACGTTCGGACGTCCGGGACAGGGGAGGTCGCACCTTAGTGGGTCACAGTGACCCGGGACGCGG
MCK promoter

GCTTCTGGCCTGGCTTTGAGTCTGAATGGCCCCCCTGGGCCCGGCCTCGTGTCCCCCCCACTGCCATCAAGGAGGGAAAACCCGCTAAGC 990
CGAAGACCGGACCGAAACTCAGACTTACCGGGGGGACCCGGGCCGGAGCACAGGGGGGGTGACGGTAGTTCCTCCCTTTTGGGCGATTCG
MCK promoter

ACAGGCATCAGGGATCAGGCTGCCCAGCTCCCACCTCTGCCCGGGTCACAGGCTCCCTGTAGCCGGGTGACAGTAGGCAAATCACGCAGC 1080
TGTCCGTAGTCCCTAGTCCGACGGGTCGAGGGTGGAGACGGGCCCAGTGTCCGAGGGACATCGGCCCACTGTCATCCGTTTAGTGCGTCG
MCK promoter

CTCTCTGGGCCACTATTTCCTCCTCTGGAGAACCAGACACTTGGTCCTTCTGGGATGATGGCAGGGTTTCCCCAGAAGCAGGGCTCAGGA 1170
GAGAGACCCGGTGATAAAGGAGGAGACCTCTTGGTCTGTGAACCAGGAAGACCCTACTACCGTCCCAAAGGGGTCTTCGTCCCGAGTCCT
MCK promoter

```
CTTTGCTGGGGTCAAGGCCACCCTGGGGGCCAAGGAGAGACTGGCGGCCTAGCGGAGGGCCAGGGGAGGGTGGTTTCTACGTGCCTGGGA   1260
GAAACGACCCCAGTTCCGGTGGGACCCCCGGTTCCTCTCTGACCGCCGGATCGCCTCCCGGTCCCCTCCCACCAAAGATGCACGGACCCT
                                        MCK promoter

CAGCCTCTGACACAGTCCCGTGGCCCCGGCGGGGGGCCAGCTGTCCCCGCCAGCCCGACTCAGCACTTGGTCTGGGGACCAGCTTGGTTT   1350
GTCGGAGACTGTGTCAGGGCACCGGGGCCGCCCCCCGGTCGACAGGGGCGGTCGGGCTGAGTCGTGAACCAGACCCCTGGTCGAACCAAA
                                        MCK promoter

GGGGGTGGGGGGTGGGCCCAGCCCCTGGGGCGGCCCATACAAGGCCATGGGGCTGGGCGCAAGGCACGCCTGGGTTCAGGGTGGGCACGG   1440
CCCCCACCCCCCACCCGGGTCGGGGACCCCGCCGGGTATGTTCCGGTACCCCGACCCGCGTTCCGTGCGGACCCAAGTCCCACCCGTGCC
                                        MCK promoter

TGCCCAGGCAGCGAAGCGAGAGCGCAGCTGCCCTCCACCCCCCTCCTGGCCAGCGGCCCCTCCTGACCAATAGCACAACCTGGGCCCCCC   1530
ACGGGTCCGTCGCTTCGCTCTCGCGTCGACGGGAGGTGGGGGGAGGACCGGTCGCCGGGGAGGACTGGTTATCGTGTTGGACCCGGGGGG
                                        MCK promoter

                                  SalI
CTATAAAGGCCAGGGCTGCAGTCCTGTCCTTTGTCGACATGGGTGAAACTCTGGGAGATTCTCTTATTGACCCAGAAAGCGATGCCTTC   1620
GATATTTTCCGGTCCCGACGTCAGGACAGGAAACAGCTGTACCCACTTTGAGACCCTCTAAGAGAATAACTGGGTCTTTCGCTACGGAAG
          MCK promoter                               PPARγ

GACACGCTGTCTGCAAACATTTCACAAGAGGTGACCATGGTTGACACCGAGATGCCGTTTTGGCCCACCAACTTTGGGATCAGCTCTGTG   1710
CTGTGCGACAGACGTTTGTAAAGTGTTCTCCACTGGTACCAACTGTGGCTCTACGGCAAAACCGGGTGGTTGAAACCCTAGTCGAGACAC
                                        PPARγ

GACCTGTCGGTGATGGACGACCACTCCCACTCCTTTGACATCAAGCCCTTCACCACTGTTGATTTCTCCAGCATTTCCACTCCACACTAT   1800
CTGGACAGCCACTACCTGCTGGTGAGGGTGAGGAAACTGTAGTTCGGGAAGTGGTGACAACTAAAGAGGTCGTAAAGGTGAGGTGTGATA
                                        PPARγ

GAAGACATCCCATTCCCGAGAGCTGATCCAATGGTTGCAGATTATAAGTACGACCTGAAGCTCCAGGACTACCAAAGTGCCATCAAAGTG   1890
CTTCTGTAGGGTAAGGGCTCTCGACTAGGTTACCAACGTCTAATATTCATGCTGGACTTCGAGGTCCTGATGGTTTCACGGTAGTTTCAC
                                        PPARγ

GAGCCTGTGTCCCCGCCTTATTATTCTGAAAAGACTCAGCTGTACAACAAACCTCACGAAGAGCCTTCCAACTCCCTCATGGCAATTGAA   1980
CTCGGACACAGGGGCGGAATAATAAGACTTTTCTGAGTCGACATGTTGTTTGGAGTGCTTCTCGGAAGGTTGAGGGAGTACCGTTAACTT
                                        PPARγ

TGCCGTGTCTGTGGGGATAAAGCCTCGGGGTTCCACTATGGAGTTCATGCTTGTGAAGGATGCAAGGGTTTCTTCCGGAGGACTATCAGA   2070
ACGGCACAGACACCCCTATTTCGGAGCCCCAAGGTGATACCTCAAGTACGAACACTTCCTACGTTCCCAAAGAAGGCCTCCTGATAGTCT
                                        PPARγ

TTGAAGCTTATTTATGATAGGTGCGATCTTAACTGTCGGATCCACAAAAAAAGTAGAAATAAATGTCAGTACTGTCGGTTTCAGAAATGC   2160
AACTTCGAATAAATACTATCCACGCTAGAATTGACAGCCTAGGTGTTTTTTTCATCTTTATTTACAGTCATGACAGCCAAAGTCTTTACG
                                        PPARγ

CTTGCTGTGGGGATGTCTCATAACGCCATCAGGTTTGGGCGGATGCCACAGGCTGAGAAGGAGAAGCTGTTGGCAGAGATCTCCAGTGAT   2250
GAACGACACCCCTACAGAGTATTGCGGTAGTCCAAACCCGCCTACGGTGTCCGACTCTTCCTCTTCGACAACCGTCTCTAGAGGTCACTA
                                        PPARγ

ATCGACCAGCTGAACCCAGAGTCTGCTGACCTCCGCGCCCTGGCAAAGCACTTGTATGACTCATACATAAAGTCCTTCCCGCTGACCAAA   2340
TAGCTGGTCGACTTGGGTCTCAGACGACTGGAGGCGCGGGACCGTTTCGTGAACATACTGAGTATGTATTTCAGGAAGGGCGACTGGTTT
                                        PPARγ
```

EcoRI

GCAAAGGCGAGGGCGATCTTGACAGGAAAGACCACAGACAAATCACCCTTTGTCATCTATGACATGAATTCCTTAATGATGGGAGAAGAT 2430
CGTTTCCGCTCCCGCTAGAACTGTCCTTTCTGGTGTCTGTTTAGTGGGAAACAGTAGATACTGTACTTAAGGAATTACTACCCTCTTCTA
PPARγ

AAAATCAAGTTCAAACACATCACCCCCCTGCAGGAGCAGAGCAAAGAGGTGGCCATTCGCATCTTTCAGGGGTGTCAGTTTCGCTCCGTG 2520
TTTTAGTTCAAGTTTGTGTAGTGGGGGGACGTCCTCGTCTCGTTTCTCCACCGGTAAGCGTAGAAAGTCCCCACAGTCAAAGCGAGGCAC
PPARγ

GAGGCCGTGCAGGAGATCACAGAGTATGCCAAGAACATCCCTGGGTTTGTAAACCTTGACCTGAATGACCAAGTAACTCTCCTAAAGTAT 2610
CTCCGGCACGTCCTCTAGTGTCTCATACGGTTCTTGTAGGGACCCAAACATTTGGAACTGGACTTACTGGTTCATTGAGAGGATTTCATA
PPARγ

GGCGTCCACGAGATCATTTACACCATGCTGGCCTCCTTGATGAATAAAGACGGGGTCCTCATCTCCGAGGGCCAAGGATTCATGACAAGG 2700
CCGCAGGTGCTCTAGTAAATGTGGTACGACCGGAGGAACTACTTATTTCTGCCCCAGGAGTAGAGGCTCCCGGTTCCTAAGTACTGTTCC
PPARγ

GAGTTTCTCAAGAGCCTGAGAAAGCCCTTTGGTGACTTTATGGAGCCCAAGTTCGAGTTTGCTGTGAAGTTCAACGCACTGGAATTAGAT 2790
CTCAAAGAGTTCTCGGACTCTTTCGGGAAACCACTGAAATACCTCGGGTTCAAGCTCAAACGACACTTCAAGTTGCGTGACCTTAATCTA
PPARγ

GACAGCGACCTGGCGATATTTATAGCTGTCATTATTCTCAGTGGAGACCGCCCAGGTTTGCTGAATGTGAAGCCCATCGAGGACATTCAA 2880
CTGTCGCTGGACCGCTATAAATATCGACAGTAATAAGAGTCACCTCTGGCGGGTCCAAACGACTTACACTTCGGGTAGCTCCTGTAAGTT
PPARγ

GACAATTTGCTGCAAGCCTTGGAGCTGCAGCTCAAGTTGAACCACCCAGAGTCCTCCCAGCTCTTTGCTAAGCTGCTCCAGAAAATGACA 2970
CTGTTAAACGACGTTCGGAACCTCGACGTCGAGTTCAACTTGGTGGGTCTCAGGAGGGTCGAGAAACGATTCGACGAGGTCTTTTACTGT
PPARγ

GACCTCAGACAGATTGTGACAGAGCACGTGCAGCTGCTGCAAGTAATAAAGAAGACAGAGACGGACATGAGCCTTCACCCACTCCTACAG 3060
CTGGAGTCTGTCTAACACTGTCTCGTGCACGTCGACGACGTTCATTATTTCTTCTGTCTCTGCCTGTACTCGGAAGTGGGTGAGGATGTC
PPARγ

NotI

GAAATATACAAGGACTTGTACTAGGCGGCCGCGACTCTAGATCATAATCAGCCATACCACATTTGTAGAGGTTTTACTTGCTTTAAAAAA 3150
CTTTATATGTTCCTGAACATGATCCGCCGGCGCTGAGATCTAGTATTAGTCGGTATGGTGTAAACATCTCCAAAATGAACGAAATTTTTT
PPARγ

CCTCCCACACCTCCCCCTGAACCTGAAACATAAAATGAATGCAATTGTTGTTGTTAACTTGTTTATTGCAGCTTATAATGGTTACAAATA 3240
GGAGGGTGTGGAGGGGGACTTGGACTTTGTATTTTACTTACGTTAACAACAACAATTGAACAAATAACGTCGAATATTACCAATGTTTAT
SV40 poly(A) signal

XhoI

AAGCAATAGCATCACAAATTTCACAAATAAAGCATTTTTTTCACTGCATTCTAGTTGTGGTTTGTCCAAACTCATCAATGTATCTCGAGA 3330
TTCGTTATCGTAGTGTTTAAAGTGTTTATTTCGTAAAAAAAGTGACGTAAGATCAACACCAAACAGGTTTGAGTAGTTACATAGAGCTCT
SV40 poly(A) signal
loxP

SalI

TAACTTCGTATAATGTATGCTATACGAAGTTATCTAGAGTCGACGGGCGGGATTCTTTTGCCCTGGCTTAACCTGATTCTTGGGCGTTGT 3420
ATTGAAGCATATTACATACGATATGCTTCAATAGATCTCAGCTGCCCGCCCTAAGAAAACGGGACCGAATTGGACTAAGAACCCGCAACA
loxP HA-R

CCTGCAGGGGATTGAGCAGGTGTACGAGGACGAGCCCAATTTCTCTATATTCCCACAGTCTTGAGTTTGTGTCACAAAATAATTATAGTG 3510
GGACGTCCCCTAACTCGTCCACATGCTCCTGCTCGGGTTAAAGAGATATAAGGGTGTCAGAACTCAAACACAGTGTTTTATTAATATCAC
HA-R

GGGTGGAGATGGGAAATGAGTCCAGGCAACACCTAAGCCTGATTTTATGCATTGAGACTGCGTGTTATTACTAAAGATCTTTGTGTCGCA 3600
CCCACCTCTACCCTTTACTCAGGTCCGTTGTGGATTCGGACTAAAATACGTAACTCTGACGCACAATAATGATTTCTAGAAACACAGCGT
HA-R

ATTTCCTGATGAAGGGAGATAGGTTAAAAAGCACGGATCTACTGAGTTTTACAGTCATCCCATTTGTAGACTTTTGCTACACCACCAAAG 3690
TAAAGGACTACTTCCCTCTATCCAATTTTTCGTGCCTAGATGACTCAAAATGTCAGTAGGGTAAACATCTGAAAACGATGTGGTGGTTTC
HA-R

TATAGCATCTGAGATTAAATATTAATCTCCAAACCTTAGGCCCCCTCACTTGCATCCTTACGGTCAGATAACTCTCACTCATACTTTAAG 3780
ATATCGTAGACTCTAATTTATAATTAGAGGTTTGGAATCCGGGGGAGTGAACGTAGGAATGCCAGTCTATTGAGAGTGAGTATGAAATTC
HA-R

***End*** (3845)

CCCATTTTGTTTGTTGTACTTGCTCATCCAGTCCCAGACATAAACATTGGCAGGGTGAGGCGGGC 3′
3845
GGGTAAAACAAACAACATGAACGAGTAGGTCAGGGTCTGTATTTGTAACCGTCCCACTCCGCCCG 5′
HA-R
